# Disruption of sphingolipid metabolism promotes tau seeding through endolysosomal membrane rigidification and rupture

**DOI:** 10.1101/2024.10.24.619578

**Authors:** Jessica Tittelmeier, Carl Alexander Sandhof, Nicole Martin, Deike El-Kabarity, Soki-Bradel Ngonza-Nito, Ronald Melki, Carmen Nussbaum-Krammer

**Affiliations:** Chair of Neuroanatomy, Institute of Anatomy, Faculty of Medicine, Ludwig-Maximilians University of Munich (LMU), Pettenkoferstrasse 11, 80336 Munich, Germany; Center for Molecular Biology of Heidelberg University (ZMBH) and German Cancer Research Center (DKFZ), DKFZ-ZMBH Alliance, Im Neuenheimer Feld 282, 69120 Heidelberg, Germany; Chemical Neurobiology Laboratory, Center for Genomic Medicine, Departments of Neurology and Psychiatry, Massachusetts General Hospital and Harvard Medical School, Boston, MA 02114, USA; Institute Francois Jacob (MIRCen), CEA, and Laboratory of Neurodegenerative Diseases, CNRS, Fontenay-Aux-Roses, France

**Keywords:** Alzheimer’s disease, tau propagation, sphingolipid metabolism, endolysosomal system, lipid homeostasis, membrane fluidity

## Abstract

Endolysosomal dysfunction is a hallmark of Alzheimer’s disease (AD) and related tauopathies, yet underlying mechanisms remain poorly understood. This study investigates the role of sphingolipid metabolism in maintaining endolysosomal membrane integrity and its impact on tau aggregation and toxicity in *Caenorhabditis elegans* and human cell culture models. Fluorescence recovery after photobleaching and C-Laurdan dye imaging revealed that silencing sphingolipid metabolism genes reduced endolysosomal vesicle membrane fluidity, increasing their rupture. The accumulation of aggregated tau in endolysosomal vesicles further aggravated endomembrane rigidification and damage and promoted seeded tau aggregation, potentially by facilitating the escape of tau seeds from the endolysosomal system. Supplementation with unsaturated fatty acids improved membrane fluidity, suppressing endolysosomal rupture and seeded tau aggregation in cell models, and alleviating tau-associated neurotoxicity in *C. elegans*. Together, this study provides mechanistic insight into how perturbation of sphingolipid metabolism promotes endolysosomal membrane damage and contributes to the escape of aggregated tau from this compartment, suggesting that restoration of membrane fluidity may represent a strategy to limit tau propagation and toxicity.

## Introduction

The gradual accumulation of microtubule-associated protein tau (MAPT/tau) aggregates is a hallmark of Alzheimer’s disease (AD) and related tauopathies. Misfolded tau species exhibit prion-like behavior by self-replicating and spreading from cell to cell^1,2^. This contributes to the progression of pathology and neurotoxicity, eventually culminating in widespread neuronal dysfunction and degeneration^3^. At the molecular level, tau aggregates replicate by templating the conversion of native tau into an amyloid conformation, promoting its accumulation into fibrillar aggregates. For this to occur at the cellular level, seeding-competent tau species (or tau “seeds”) must be released from a donor cell and taken up into the cytosol of the neighboring receiving cell in order to come into direct contact with the cytosolic native tau protein^4–6^.

In this process, the autophagy-lysosomal pathway (ALP), which is an important clearance route for tau^7^, appears to play a central role. Studies have demonstrated impaired ALP function in the brains of tauopathy patients as well as in animal and cell models, showing that the accumulation of abnormal autolysosomal and endolysosomal vesicles correlating with neuronal toxicity^8–11^. Inhibition of autophagic clearance of tau increases its secretion and spreading^12^. In addition, the accumulation of misfolded tau within endolysosomes leads to a destabilization and rupture of these vesicles^13–18^. While intact endolysosomes normally restrict tau seeds from reaching cytosolic monomers, membrane rupture allows their escape. Recent studies highlight this escape as a critical rate-limiting step in seeded tau propagation^13–18^. Endolysosomal damage not only promotes the propagation of pathological aggregates, but also causes the release of hydrolytic enzymes into the cytosol, leading to cellular damage and death^19^. Although recent efforts to identify pathways involved in endolysosomal damage and repair have intensified, many aspects of these processes remain unknown.

To identify cellular factors that are critical for the integrity of endolysosomal vesicles, we recently performed an unbiased genome-wide RNA interference (RNAi) screen in *C. elegans*^20^. One of the pathways identified was sphingolipid (SL) metabolism. SLs constitute a large and diverse class of lipids that are involved in various physiological processes^21, 22^. They consist of two main building blocks, a long-chain base with a serine backbone and an attached long acyl chain. This characteristic chemical structure mediates unique biophysical properties. SL biosynthesis and degradation rely on a specialized enzymatic machinery distinct from that of other lipids^23^ (Figure 1A). Functionally, SLs, along with glycerolipids and sterols, serve as structural components of cell membranes, contributing to the stability and fluidity of membranes and to the organization of microdomains such as lipid rafts. Beyond their structural role, SLs serve as bioactive molecules participating in cellular signaling pathways that control cell growth, differentiation, apoptosis, and intercellular communication^24^. Abnormalities in SL metabolism have been observed during aging and in neurodegenerative conditions, including AD, underscoring their potential significance in the pathogenesis of these disorders^25, 26^. Interestingly, recent evidence suggests that sphingolipid accumulation disturbs the endolysosomal pathway and induces or potentiates endolysosomal membrane rupture^27, 28^; however, the mechanisms underlying membrane destabilization remain unclear.

**Figure 1.**
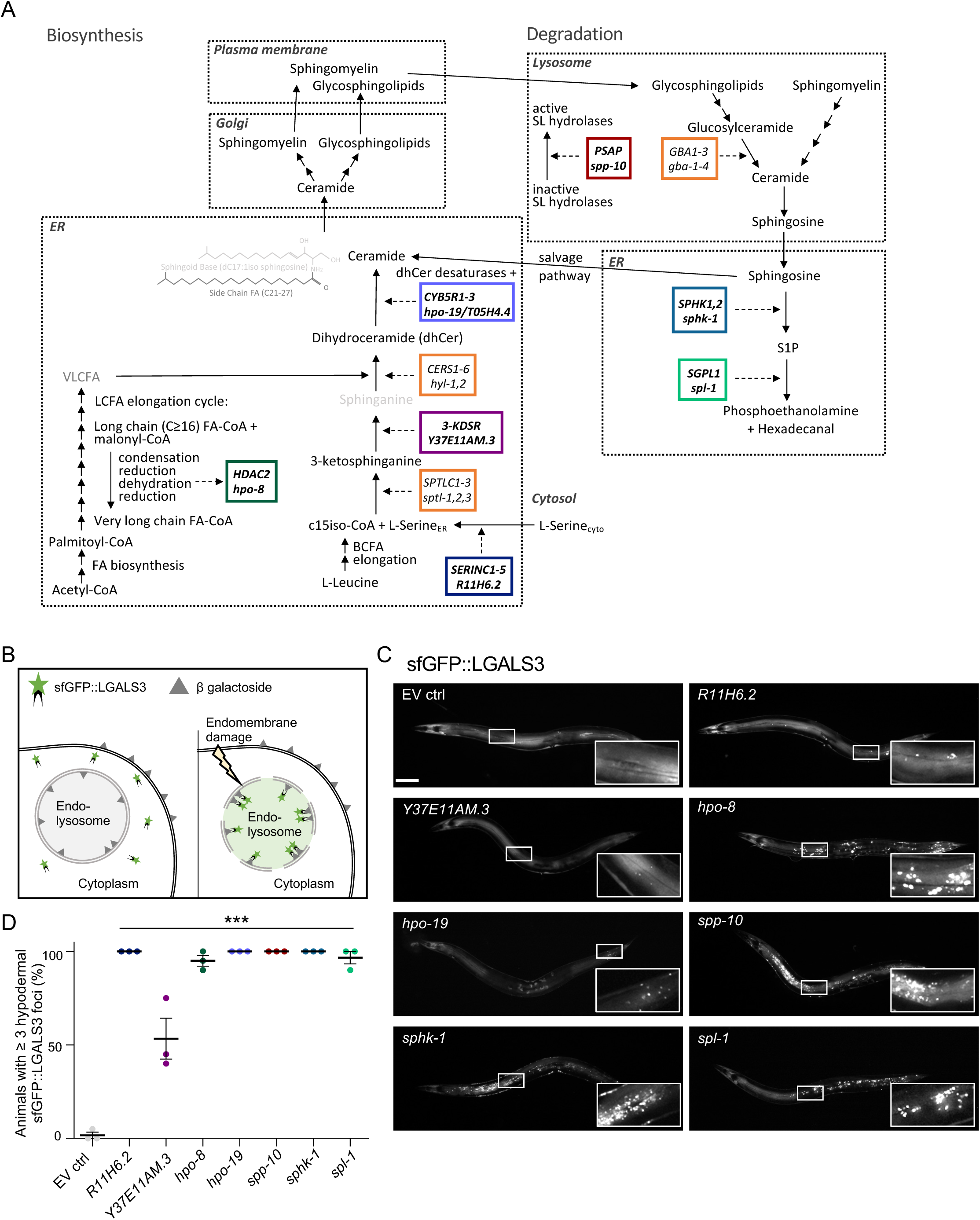
Knockdown of sphingolipid metabolism genes promotes endolysosomal vesicle rupture. **(A)** General overview of sphingolipid metabolism with a particular focus on the genes identified in the genome-wide screen. Sphingolipids constitute a group of amphipathic lipids featuring a polar head group and a sphingoid base backbone that is N-acetylated with a (very) long-chain fatty acid ((V)LCFA) side chain. In contrast to mammals, where the sphingoid base is conventionally derived from palmitic acid and serine, *C. elegans* sphingolipids usually contain a characteristic C17iso branched chain sphingoid base^75–77^. Its synthesis involves the branched chain FA (BCFA) elongation pathway to yield C15iso-CoA, which then condenses with L-serine to form 3-ketosphinganine. This reaction is catalyzed by serine palmitoyltransferase (encoded by *sptl-1*, -*2*, and *-3*). The serine incorporator (SERINC) protein family (encoded by *R11H6.2*) is believed to assist in the incorporation of L-serine into specific membranes. 3-ketodihydrosphingosine reductase (KDSR, in *C. elegans* predicted to be encoded by *Y37E11AM.3*) then catalyzes the reduction of 3-keto sphinganine to sphinganine. The (V)LCFA side chain is primarily comprised of a straight saturated FA chain, ranging from 20-26 carbon atoms in length, with or without hydroxylation^30, 78^. It can also be derived from BCFAs, such as C15iso and C17iso^75, 76^. However, most of the side chain FA moieties originate from palmitoyl-CoA via the canonical de novo FA biosynthesis pathway, involving the sequential addition of C2 moieties from malonyl-CoA through the LCFA elongation cycle^75, 76^. Each elongation cycle comprises four reactions (condensation, reduction, dehydration, reduction), with the third reaction requiring very-long-chain (3R)-3-hydroxyacyl-CoA dehydratase (encoded by *hpo-8*)^76^. Finally, ceramide synthases (encoded by *hyl-1* and *hyl-2*) catalyze the addition of various acyl side chains to the sphingoid base to yield dihydroceramide, which is then desaturated to ceramide. The latter reaction is catalyzed by dihydroceramide desaturases, which require electrons from NAD(P)H provided by cytochrome b5 reductases (encoded by *hpo-19* and *T05H4.4*). All complex sphingolipids, such as sphingomyelin and glycosphingolipids (including cerebrosides and gangliosides) originate from ceramide. Degradation of complex sphingolipids takes place in the lysosome. Essential for this process are saposins or sphingolipid activator proteins (PSAPs, encoded by *spp-10*), which serve as crucial bridges between the lipid substrate and hydrophilic hydrolases. Glucocerebrosidases (encoded by *gba-1*, *gba-2*, *gba-3*, and *gba-4*) hydrolyse glucosylceramide into ceramide and glucose. Sphingosine may be either recycled and metabolized back into ceramide or phosphorylated by sphingosine kinase (encoded by *sphk-1*) to generate sphingosine-1-phosphate (S1P). S1P lyase (encoded by *spl-1*) irreversibly cleaves S1P into phospho-ethanolamine and hexadecenal. The *C. elegans* genes identified in the primary screen, along with their human orthologs, are framed with color. Genes identified in subsequent co-RNAi experiments and their human orthologs are framed in gray. **(B)** Schematic of lysosomal rupture detected by the galectin puncta assay. **(C)** Widefield fluorescence images of day 5 (second day of adulthood) animals expressing F3ΔK281::mCherry in touch receptor neurons and hypodermal sfGFP::LGALS3 and RNAi mediated KD of indicated sphingolipid metabolism genes. Numerous foci are visible indicating lysosomal rupture. Zoomed in image is indicated in overview image by a white box. Scale bar 100 µm. **(D)** Mean percentage of day 5 (second day of adulthood) animals positive for lysosomal rupture (defined as three or more sfGFP::LGALS3 foci in the hypodermis). Data shown as means of three technical replicate plates with 17-30 animals per plate ± SEM. Statistical analysis comparing RNAi conditions to the empty vector control (EV ctrl) was done using one-way ANOVA with Dunnett’s post-hoc test. *** = p < 0.001.

Here we investigated the role of SL metabolism in maintaining endolysosomal vesicle integrity. Our results revealed that silencing of SL metabolism genes led to a marked reduction in membrane fluidity, particularly in membranes of endolysosomal vesicles, thereby increasing their fragility and susceptibility to rupture. The accumulation of aggregated tau had an additive effect and led to a further reduction in endomembrane fluidity, which aggravated the damage to the endolysosomal compartment. Increased membrane rigidity also facilitated seeded tau aggregation. Conversely, improving membrane fluidity through supplementation with polyunsaturated fatty acids (PUFAs) counteracted tau propagation in cells and tau-associated neurotoxicity in *C. elegans*.

This study highlights the interplay between lipid metabolism and proteostasis and provides insights into how perturbations in sphingolipid metabolism contribute to endolysosomal membrane dysfunction and tau-associated phenotypes. In addition, our data offer a potential mechanistic explanation for the beneficial effects of PUFA-rich diets or PUFA supplementation reported in AD patients and suggest that restoring membrane fluidity may help mitigate tau toxicity and disease progression.

## Results

### Disruption of sphingolipid metabolism induces endolysosomal rupture

Alterations in SL metabolism are increasingly recognized in aging and neurodegenerative diseases, including AD; however, it remains unclear whether these changes actively contribute to disease pathology or merely reflect downstream consequences of neurodegeneration. In our previously published unbiased genome-wide RNAi screen in *C. elegans*, we identified SL metabolism as one of the pathways whose disruption compromises endolysosomal membrane integrity^20^. Here, we focused on these SL-related hits to better understand how perturbation of SL metabolism promotes endolysosomal rupture and whether it affects tau-related phenotypes.

To first confirm the SL-related hits under the same assay conditions in which they were originally identified, we used the *C. elegans* reporter strain from our previously published screen^20^ (Table S1). In this strain, endolysosomal membrane damage is monitored in the hypodermis by expression of human galectin-3 fused to superfolder-GFP (sfGFP::LGALS3). The animals also express an aggregation-prone tau fragment fused to mCherry (F3ΔK281::mCherry) in touch receptor neurons, which is transmitted to the hypodermis, as described previously^20^. Under steady-state conditions, sfGFP::LGALS3 remains diffusely distributed throughout the cytosol. Upon endolysosomal damage, luminal β-galactosides become exposed and recruit sfGFP::LGALS3 into visible puncta, providing a sensitive readout of vesicle rupture^20,29^ (Figure 1B).

Using this reporter system, we confirmed that knockdown (KD) of selected SL-related hits increased sfGFP::LGALS3 foci formation. These hits included the serine incorporator *R11H6.2*, the 3-ketodihydrosphingosine reductase *Y37E11AM.3,* the hydroxyacyl-CoA dehydratase *hpo-8*, the cytochrome b5 reductases *hpo-19* and *T05H4.4,* the saposin *spp-10*, the sphingosine kinase *sphk-1*, and the sphingosine-1-phosphate lyase *spl-1* (Figure 1C, D). As genetic validation independent of RNAi, we tested an available *sphk-1* mutant strain, which also showed a robust increase in hypodermal sfGFP::LGALS3 foci (Figure S1A). These data support the conclusion that genetic perturbation of sphingolipid metabolism compromises endolysosomal integrity.

The screen identified distinct steps in sphingolipid metabolism catalyzed by non-redundant genes or targeted by a single RNAi clone, as in the case of *hpo-19*/*T05H4.4* which are both depleted by *hpo-19* RNAi (Figure 1A). However, other steps in this pathway, which are mediated by two or more proteins may have been missed in the screen. Indeed, simultaneous KD of selected redundant genes using co-RNAi revealed additional genes functioning in *de novo* sphingolipid biosynthesis that also induced endolysosomal rupture, including the serine palmitoyltransferases *sptl-1* and -*3* and the ceramide synthases *hyl-1* and -*2* (Figure S1B). In addition, the co-KD of all four glucocerebrosidases *gba-1-4* or only *gba-2-4*, also triggered endolysosomal rupture (Figure S1C). Together, these results indicate that perturbing SL metabolism at multiple steps can compromise endolysosomal integrity, highlighting the tight metabolic balance required to maintain the stability of the endolysosomal limiting membrane. Because KD efficiency was not assessed for the individual RNAi clones or co-RNAi combinations, these experiments do not allow comparison of relative RNAi strength or inference of the relative importance of individual genes. Thus, the conclusions drawn from these RNAi experiments are qualitative: specific single or combined KDs can promote endolysosomal rupture, whereas the absence of a detectable phenotype after RNAi cannot exclude gene involvement, as KD may have been insufficient.

### Disruption of sphingolipid metabolism reduces endolysosomal membrane fluidity in *C. elegans*

SLs are important components of eukaryotic cell membranes and genetic modulation of SL metabolism likely affects the lipid composition and biophysical properties of membranes. Similar to their mammalian isoforms, the N-acyl chains of *C. elegans* SLs contain predominantly long and saturated fatty acid chains^30, 31^, which tend to pack tightly within the lipid bilayer. As such, a higher proportion of SLs generally reduces membrane fluidity, leading to increased order, rigidity, or viscosity^32^. Consequently, KD of the hits involved in SL degradation should lead to SL accumulation and reduced membrane fluidity, while KD of the hits involved in SL biosynthesis should have the opposite effect.

To evaluate the membrane fluidity of endolysosomal membranes, fluorescence recovery after photobleaching (FRAP) was employed on *C. elegans* that express the Lysine/Arginine Transporter 1 (LAAT-1) tagged with mCherry, which localizes to the lysosomal membrane^33^. FRAP enables the assessment of the lateral mobility of LAAT-1 within the lysosomal membrane, thereby indirectly measuring its fluidity. We focused our analysis on the hypodermis, where we expressed the sfGFP::LGALS3 reporter and observed endolysosomal rupture following KD of the SL metabolism genes (Figure 1C, D). Interestingly, KD of all of our hits significantly increased the time required to recover half of the maximum fluorescence intensity (t_half_), indicating less fluid lysosomal membrane in the hypodermis (Figure 2A-C, E, F). With some gene KDs, we also observed a decrease in the maximal recovered signal after bleaching (Figure 2D, G). The decrease in lysosomal membrane fluidity was not specific to the hypodermis as the KD of *spl-1* also increased the t_half_ of LAAT-1::mCherry in intestinal lysosomes (Figure S2A-D). To discern whether the detrimental effect of KD on membrane fluidity is confined to lysosomal membranes or extends to other cellular membranes, we utilized a *C. elegans* strain expressing a plasma membrane-anchored GFP^34^. Intriguingly, this reporter showed no change in the t_half_ with any KD and only *spl-1* KD led to decrease in the maximal recovered signal (Figure 2H-K, Figure S2E-G). Hence, the endolysosomal membrane seems to be particularly sensitive to a disruption of SL metabolism in contrast to the plasma membrane. Perturbing genes involved in both SL synthesis and degradation reduces the fluidity of the endolysosomal membrane and makes it more susceptible to rupture.

**Figure 2.**
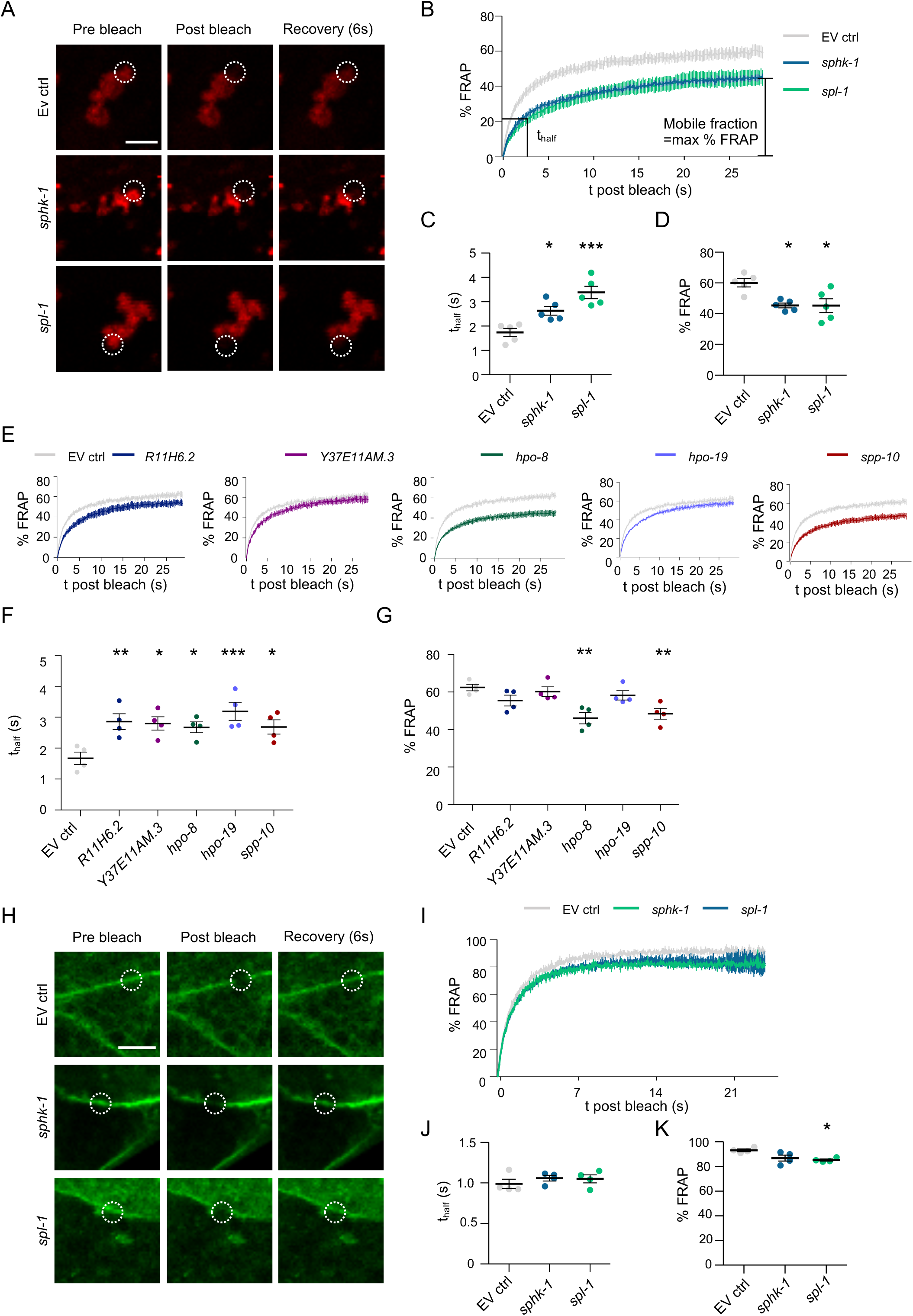
Knockdown of genes involved in sphingolipid metabolism decreases lysosomal membrane fluidity. **(A)** Representative confocal single plane images from a FRAP experiment in day 5 (second day of adulthood) animals expressing a mCherry tagged Lysosomal Lysine/Arginine Transporter 1 (LAAT-1::mCherry) in the hypodermis grown either on empty vector control (EV ctrl), *spl-1* or *sphk-1* RNAi plates. Dashed circles outline the bleach spots. Scale bar = 10 µm. **(B)** Combined FRAP curves of LAAT-1::mCherry in hypodermal lysosomal membranes. Curves are normalized to the pre-bleach intensity as 100% and the first post-bleach intensity as 0%. **(C, D)** Increase (C) in the mean time until half of the maximal signal is recovered (t_half_,) and decrease (D) in the maximal % recoverable fluorescence values upon KD of *sphk-1* and *spl-1* indicate a reduction in lysosomal membrane fluidity. **(E)** FRAP curves of LAAT-1::mCherry in hypodermal lysosomal membranes of animals grown either on empty vector or the indicated RNAi plates. Curves are normalized to the pre-bleach intensity set as 100% and the first post-bleach intensity as 0%. **(F)** Mean t_half_ upon KD of sphingolipid metabolism genes. **(G)** Maximal % recoverable fluorescence values upon KD of sphingolipid metabolism genes. **(H)** Representative confocal single plane images from a FRAP experiment in animals expressing prenylated GFP for lipid membrane anchorage in the intestine. Scale bar = 5 µm. **(I)** Combined FRAP curves of prenylated GFP enriched on the intestinal plasma membrane of animals grown either on empty vector, *sphk-1* or *spl-1* RNAi plates. Curves are normalized to the pre-bleach intensity as 100% and the first post-bleach intensity as 0%. **(J, K)** Mean t_half_ (J) and maximal % recoverable fluorescence values (K). Data represented as means ± SEM of 5 - 12 FRAP measurements per condition in animals on day 5 (second day of adulthood) collected in five biological replicates. Statistical analysis comparing RNAi conditions to the empty vector control was done using one-way ANOVA with Dunnett’s post-hoc test. n.s.: not significant, * = p < 0.05, ** = p < 0.01, *** = p < 0.001.

### KD of SPHK2 increases endolysosomal membrane rigidity in human cells

Having established in *C. elegans* that disruption of SL metabolism reduces endolysosomal membrane fluidity, we next asked whether this relationship is conserved in human cells. To this end, we employed SH-SY5Y human neuroblastoma cells and assessed membrane fluidity using C-Laurdan dye (Figure 3A). This approach leverages the unique properties of C-Laurdan, which emits fluorescent light of varying wavelengths in response to changes within the phospholipid bilayer, particularly reflective of membrane fluidity^35, 36^. The shift in emission profile between liquid-disordered and liquid-ordered phases allows a quantitative assessment of the membrane order by calculating the ratiometric relationship of the fluorescence intensity recorded in two spectral channels, known as a generalized polarization (GP) value^35^ (Figure 3A). Decreasing GP values are indicative of increasingly fluid membrane while increasing GP values denote increased rigidity. Of note, we exposed the cells to C-Laurdan for an extended time of 2 hours (instead of 30 min as recommended in the original protocol), to promote internalization of the dye and thus increase the staining of endolysosomal membranes relative to the plasma membrane^35^.

**Figure 3.**
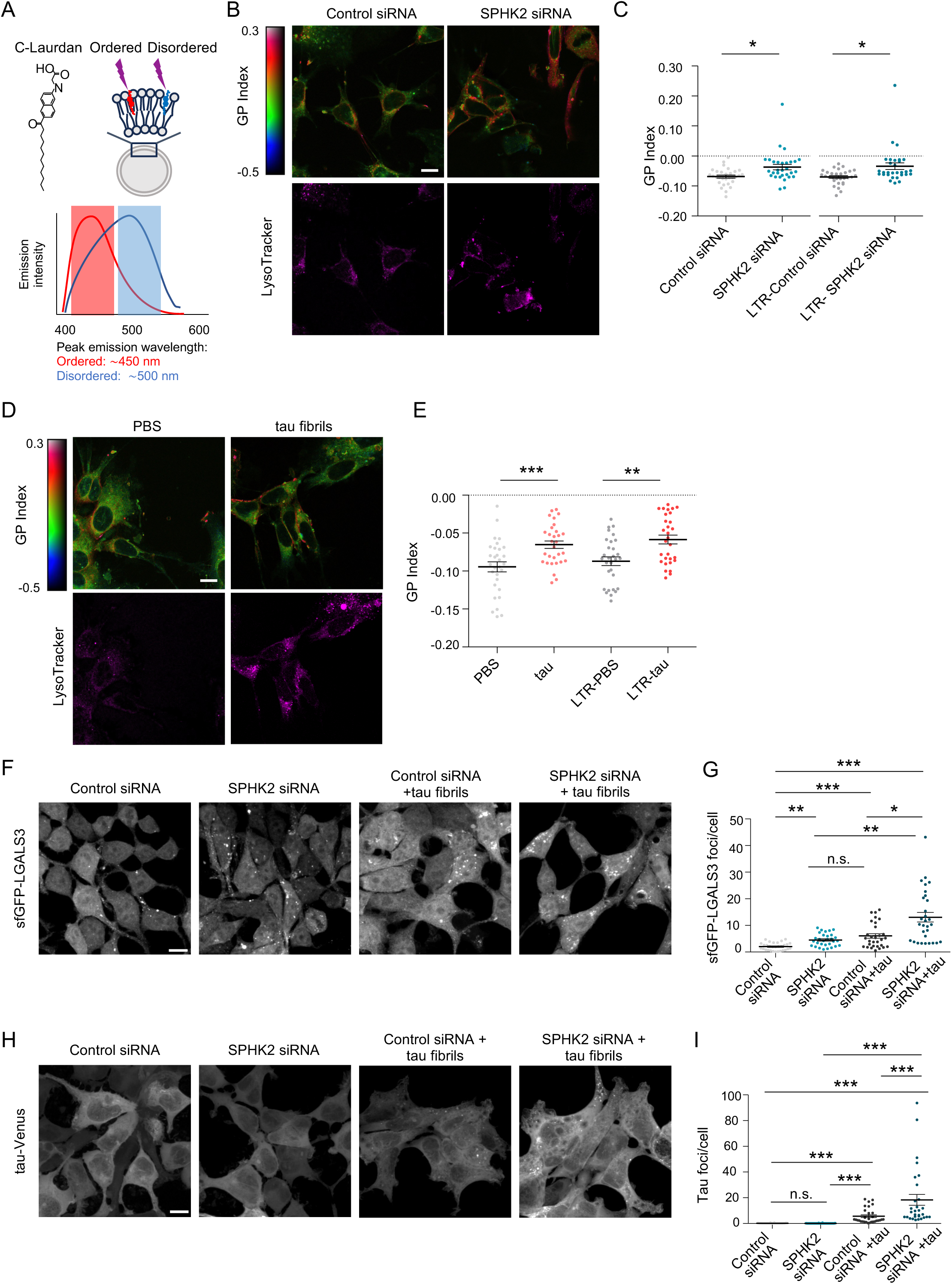
KD of SPHK2 and aggregated tau increases membrane rigidity leading to lysosomal rupture. **(A)** Scheme of the fluorescence properties of C-Laurdan. The dye is excited at 405 nm and exhibits peak emission at 450 nm (red) in ordered membrane phases and ∼500 nm in the disordered phase (blue). Two-channel acquisition is conducted in the wavelength bands indicated by shaded boxes. **(B)** Upper panels: Pseudo-colored images of SH-SY5Y cells transfected with control or SPHK2 siRNA showing the C-Laurdan GP Index at each pixel position. Lower panels show LysoTracker staining. Scale bar = 10 µm. **(C)** Quantification of GP values in SH-SY5Y cells upon transfected with control or SPHK2 siRNA. GP values were measured across the whole cell (left) or restricted to LysoTracker-positive (LTR) regions (right). Statistical analysis was conducted using a two-way mixed-model ANOVA, followed by pairwise comparisons of estimated marginal means with Sidak correction for multiple comparisons. n = 3 independent experiments, with 10 images analyzed per experiment. **(D)** Upper panel: Pseudo-colored images of SH-SY5Y cells exposed to PBS control or 1N4R tau fibrils showing the C-Laurdan GP Index at each pixel position. Lower panels show LysoTracker staining. Scale bar = 10 µm. **(E)** Quantification of GP values in SH-SY5Y cells exposed to PBS control or 1N4R tau fibrils. GP values were measured across the whole cell (left) or restricted to LysoTracker-positive (LTR) regions (right). Statistical analysis was conducted using a two-way mixed-model ANOVA followed by pairwise comparisons of estimated marginal means with Sidak correction for multiple comparisons. n = 3 independent experiments, with 10 images analyzed per experiment. **(F)** Maximum intensity projection of confocal z-stacks of HEK293T cells expressing sfGFP-LGALS3 upon treatment with control or SPHK2 siRNA, exposed to PBS control or 1N4R tau fibrils. Scale bar = 10 µm. **(G)** Quantification of sfGFP-LGALS3 foci per cell in HEK293T cells upon treatment with control or SPHK2 siRNA, exposed to PBS control or 1N4R tau fibrils. Data represent the mean number of foci per cell. Statistical analysis was done using Kruskal-Wallis with a Dunn’s post-hoc test. N = 3 independent experiments, with 10 images analyzed per experiment. **(H)** Maximum intensity projection of confocal z-stacks of a P301S tau-Venus biosensor cell line upon treatment with control or SPHK2 siRNA with and without exposure to 1N4R tau fibrils. Scale bar = 10 µm. **(I)** Quantification of tau-Venus foci upon treatment with control or SPHK2 siRNA with and without exposure to 1N4R tau fibrils. Data were analyzed by a two-way mixed-model ANOVA, followed by pairwise comparisons were performed using estimated marginal means with Sidak correction. n = 3 independent experiments, with 10 images analyzed per experiment. n.s.: not significant, * = p < 0.05, ** = p < 0.01, *** = p < 0.001.

Upon KD of *SPHK2*, one of the human homologs of *C. elegans sphk-1*, the C-Laurdan fluorescence GP Index increased significantly, indicating increased membrane rigidity (Figure 3B, C). Lipofectamine treatment alone did not alter GP values (Figure S3A, B), and SPHK2 KD was confirmed by immunoblotting after 48 h and 72 h (Figure S3C–E). To determine whether SPHK2 KD affects lysosomal membrane fluidity, we combined C-Laurdan staining with LysoTracker staining and selectively analyzed LysoTracker-positive regions. SPHK2 KD resulted in a pronounced increase in GP values within LysoTracker-positive compartments, demonstrating increased membrane rigidity at lysosomes (Figure 3C). Taken together, FRAP and C-Laurdan analyses revealed that disruption of SL metabolism leads to more ordered and rigid endolysosomal membranes in both *C. elegans* and human cells.

### Fibrillar tau and SPHK2 KD act in concert to exacerbate endolysosomal damage and seeded tau aggregation

Given that SPHK2 KD increased endolysosomal membrane rigidity and can compromise endolysosomal integrity, we next asked whether aggregated tau similarly alters membrane properties. Since fibrillar tau is known to induce endolysosomal rupture^14^, we hypothesized that tau-induced membrane damage may also involve increased membrane rigidity. Treatment of SH-SY5Y and HEK293T cells with recombinant 1N4R tau fibrils led to a marked increase in membrane rigidity, as indicated by elevated GP values (Figure 3D, E, Figure S3F, G). This increase was also observed in LysoTracker-positive compartments, indicating increased rigidity of lysosomal membranes. Importantly, this effect was specific to fibrillar tau, as monomeric tau did not alter membrane fluidity (Figure S3H, I). These findings suggest that aggregated tau, like SPHK2 KD, promotes membrane rigidification, providing a potential mechanism by which tau fibrils may contribute to endolysosomal rupture.

We therefore asked whether SPHK2 KD further enhances tau-fibril-induced endolysosomal damage. To test this, we used a previously established HEK293T cell line stably expressing sfGFP-LGALS3 and monitored endolysosomal rupture by galectin puncta formation^20,29^. Cells were treated with recombinant 1N4R tau fibrils, either alone or in combination with siRNA-mediated SPHK2 KD. Both, tau fibrils and SPHK2 KD alone induced endolysosomal rupture, as evidenced by increased sfGFP-LGALS3 puncta formation (Figure 3F, G and Figure S3J). While SPHK2 KD alone significantly increased galectin puncta above the matched control, its effect was more modest than in our previous CRISPR inhibition-based analysis^20^. This difference likely stems from the earlier readout required for the combined siRNA/tau fibril assay, when transient Lipofectamine-associated effects still increased the control background. Importantly, SPHK2 KD further enhanced tau fibril-induced sfGFP-LGALS3 foci formation, indicating that SL disruption compromises endolysosomal integrity and sensitizes endolysosomal membranes to tau fibril-induced damage in human cells.

Because endolysosomal rupture facilitates the escape of luminal contents, including tau seeds, into the cytosol, we next asked whether membrane perturbation by SPHK2 KD influences seeded tau aggregation. Using a HEK biosensor cell line expressing Venus-tagged full-length P301S mutant 0N4R tau (tau-Venus)^37, 38^, we found that SPHK2 KD alone did not induce tau aggregation, as no increase in tau-Venus foci was observed (Figure 3H, I and Figure S3K). However, upon addition of recombinant tau fibrils, SPHK2 KD significantly increased tau-Venus foci formation. Thus, under the conditions tested here, disruption of SL metabolism alone is not sufficient to initiate detectable tau aggregation. Rather, it seems to facilitate the escape of tau seeds from the endolysosomal compartment into the cytosol by compromising endolysosomal membrane integrity.

### Tau transmission sensitizes endolysosomal membranes to sphingolipid perturbations in vivo

To determine how tau transmission and SL perturbations interact to affect endolysosomal integrity in vivo, we returned to *C. elegans*. We compared animals expressing F3ΔK281::mCherry in touch receptor neurons, from where it is transmitted to the hypodermis, with matched controls expressing mCherry alone in the same neurons^20^. In both strains, sfGFP::LGALS3 is expressed in the hypodermis to monitor endolysosomal membrane damage^20^. KD of Y37E11AM.3 alone did not increase hypodermal sfGFP::LGALS3 puncta compared to EV control RNAi, whereas the presence of transmitted F3ΔK281::mCherry significantly increased galectin puncta (Figure 4A). In contrast, KD of the remaining SL-related hits resulted in nearly all animals displaying hypodermal sfGFP::LGALS3 foci in both genetic backgrounds. This suggests that strong disruption of SL metabolism is sufficient to overwhelm endolysosomal integrity independently of tau.

**Figure 4.**
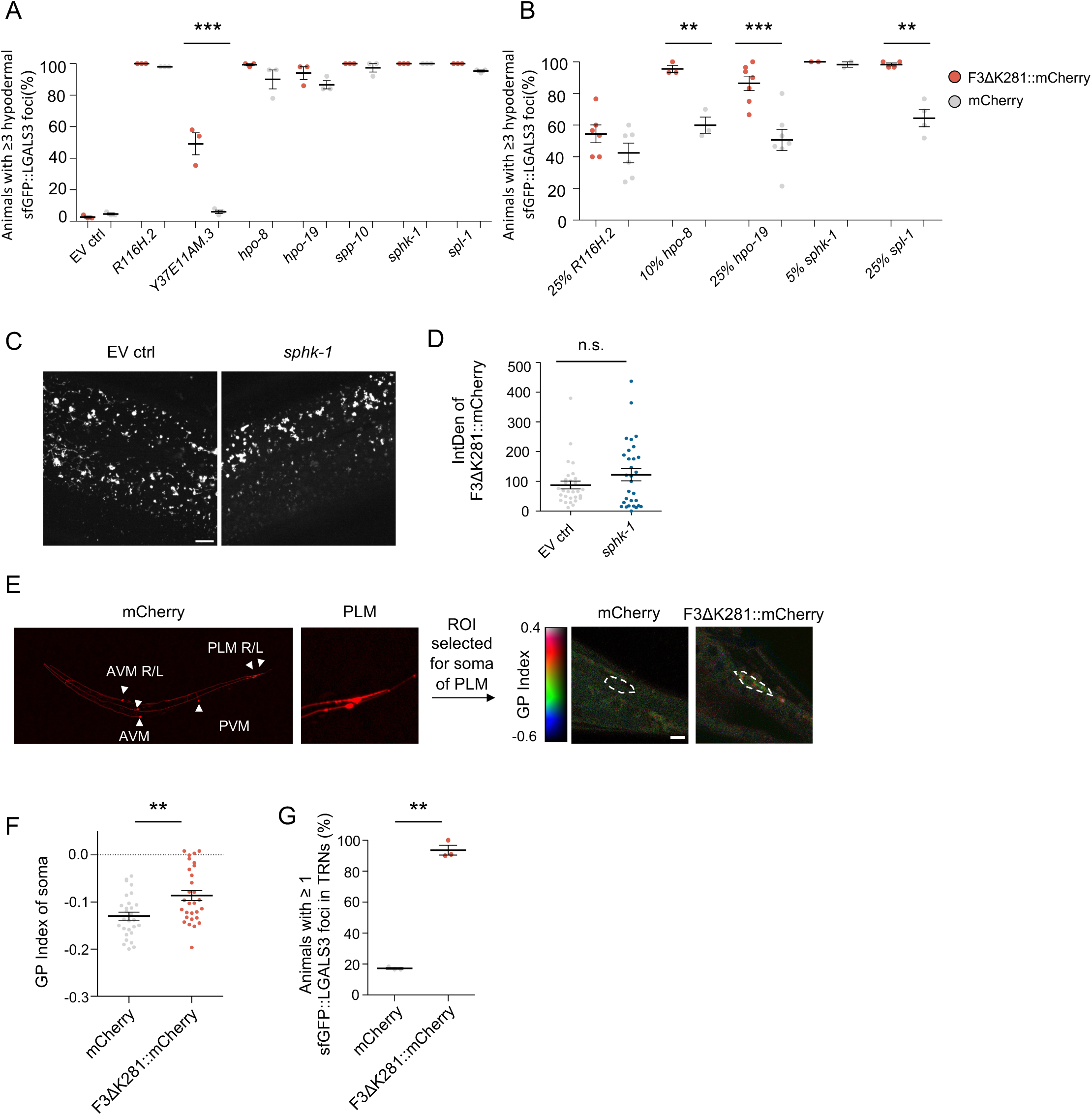
Aggregated tau promotes endolysosomal rupture. **(A, B)** Quantification of the percentage of animals with ≥ 3 hypodermal sfGFP::LGALS3 foci, upon expression of either F3ΔK281::mCherry (red) or mCherry control (grey) in touch receptor neurons and KD of the indicated genes on day 5 (second day of adulthood) (A). Quantification of endolysosomal rupture under sub-saturating RNAi conditions. RNAi cultures were diluted with the empty vector (EV) control bacteria to reduce the KD strength (B). Data represented as mean ± SEM. n = 3 - 7 independent experiments with 40 - 50 (A) or 20 - 30 (B) animals analyzed per experiment. Since even a dilution down to 5% of the *sphk-1* RNAi resulted in 100% of animals being scored as positive in two independent replicates this condition was not repeated further. Statistical analysis comparing mCherry to F3ΔK281::mCherry under individual RNAi conditions was done using two-way ANOVA with Sidaks’s post-hoc test. ** = p < 0.01, *** = p < 0.001. **(C)** Representative images of hypodermal F3ΔK281::mCherry signal in EV control or *sphk-1* RNAi treated animals on day 5 (second day of adulthood). Scale bar = 10 µm. **(D)** Quantification of hypodermal F3ΔK281::mCherry fluorescence intensity (integrated density, IntDen) in EV control or *sphk-1* RNAi treated animals on day 5 (second day of adulthood), indicating *sphk-1* KD does not alter tau transmission levels. Statistical analysis was done using Student’s t-test. n = 3 independent experiments with 35 animals analyzed in total. **(E)** C-Laurdan staining of animals expressing F3ΔK281::mCherry or mCherry in touch receptor neurons on day 4 (first day of adulthood). The mCherry signal was used to select a region of interest (ROI) around the soma of the posterior touch receptor neurons (PLM) to determine the GP value. Scale bar = 10 µm. **(F)** Quantification of GP values in animals expressing F3ΔK281::mCherry compared to mCherry on day 4 (first day of adulthood). Statistical analysis was done using a Student’s t-test. n = 4 independent experiments, with at least 28 animals analyzed in total. **(G)** Quantification of the percentage of animals expressing mCherry or F3ΔK281::mCherry in touch receptor neurons with ≥1 sfGFP::LGALS3 puncta in touch receptor neurons on day 5 (second day of adulthood). Each dot represents an independent experiment, and lines indicate the mean ± SEM. n = 3 independent experiments, with 10-14 animals per experiment. Statistical analysis was done using a Student’s t-test. n.s.: not significant, * = p < 0.05, ** = p < 0.01, *** = p < 0.001.

To determine whether tau-dependent effects were masked by the strength of RNAi, we titrated RNAi bacterial concentrations to achieve sub-saturating KD conditions. Under these conditions, KD of *spl-1, hpo-8,* and *hpo-19* induced endolysosomal damage that was significantly exacerbated in F3ΔK281::mCherry animals compared to mCherry controls (Figure 4B). In contrast, dilution of *R11H6.2* RNAi did not reveal tau-dependent differences, and *sphk-1* KD remained fully penetrant even at high dilution. Importantly, RNAi-mediated knockdown of *sphk-1* did not alter hypodermal F3ΔK281::mCherry levels, arguing that the enhanced rupture phenotype is not due to increased tau transmission (Figure 4C, D).

To assess whether tau directly alters membrane properties in vivo, we measured membrane fluidity in the somas of the posterior touch receptor (PLM) neurons using C-Laurdan dye. Expression of the tau fragment led to a decrease in membrane fluidity compared to mCherry controls (Figure 4E, F), indicating that tau promotes membrane rigidification in PLM neurons, consistent with our observations in human cells (Figure 3D, E). In addition, expression of sfGFP-LGALS3 in touch receptor neurons revealed increased galectin puncta in animals expressing F3ΔK281::mCherry compared to mCherry controls, indicating elevated endolysosomal rupture in neurons (Figure 4G). These findings suggest that perturbation of SL metabolism and tau accumulation both increase endolysosomal membrane rigidity and make vesicles more prone to rupture, with additive effects when both perturbations occur simultaneously.

### Increasing membrane fluidity reduces aggregated tau-mediated endolysosomal damage, seeded aggregation, and neurotoxicity

SL metabolism can influence cellular physiology through multiple mechanisms beyond its effects on membrane fluidity. To test more directly whether altered membrane fluidity contributes to aggregated tau-induced endolysosomal damage and propagation, we used fatty acid supplementation as an independent approach to modulate lipid packing. Saturated fatty acids such as palmitic acid (PA) promote tight lipid packing and increase membrane rigidity, whereas unsaturated fatty acids contain one or more double bonds that introduce kinks into their hydrocarbon chains, resulting in looser lipid packing and increased membrane fluidity.

We first increased membrane rigidity with PA. PA treatment increased membrane rigidity in SH-SY5Y and HEK293T cells (Figure S4A–F), exacerbated tau fibril-induced sfGFP-LGALS3 foci formation (Figure S4G,H), and enhanced tau-Venus aggregation after exposure to tau fibrils (Figure S4I,J). These findings support a model in which increased membrane rigidity promotes endolysosomal damage and facilitates tau propagation.

We next asked whether increasing membrane fluidity could counteract these effects. Treatment of SH-SY5Y and HEK293T cells with the ω-3 polyunsaturated fatty acid (PUFA) α-linolenic acid (ALA) increased membrane fluidity (Figure S5A-C) and prevented tau fibril-induced membrane rigidification (Figure 5A, B and Figure S5D). Consistent with this, LysoTracker-based analysis indicated that ALA also affected lysosome-associated membrane properties. Moreover, ALA pre-treatment reduced tau fibril-induced endolysosomal rupture (Figure 5C, D and Figure S5E).

**Figure 5.**
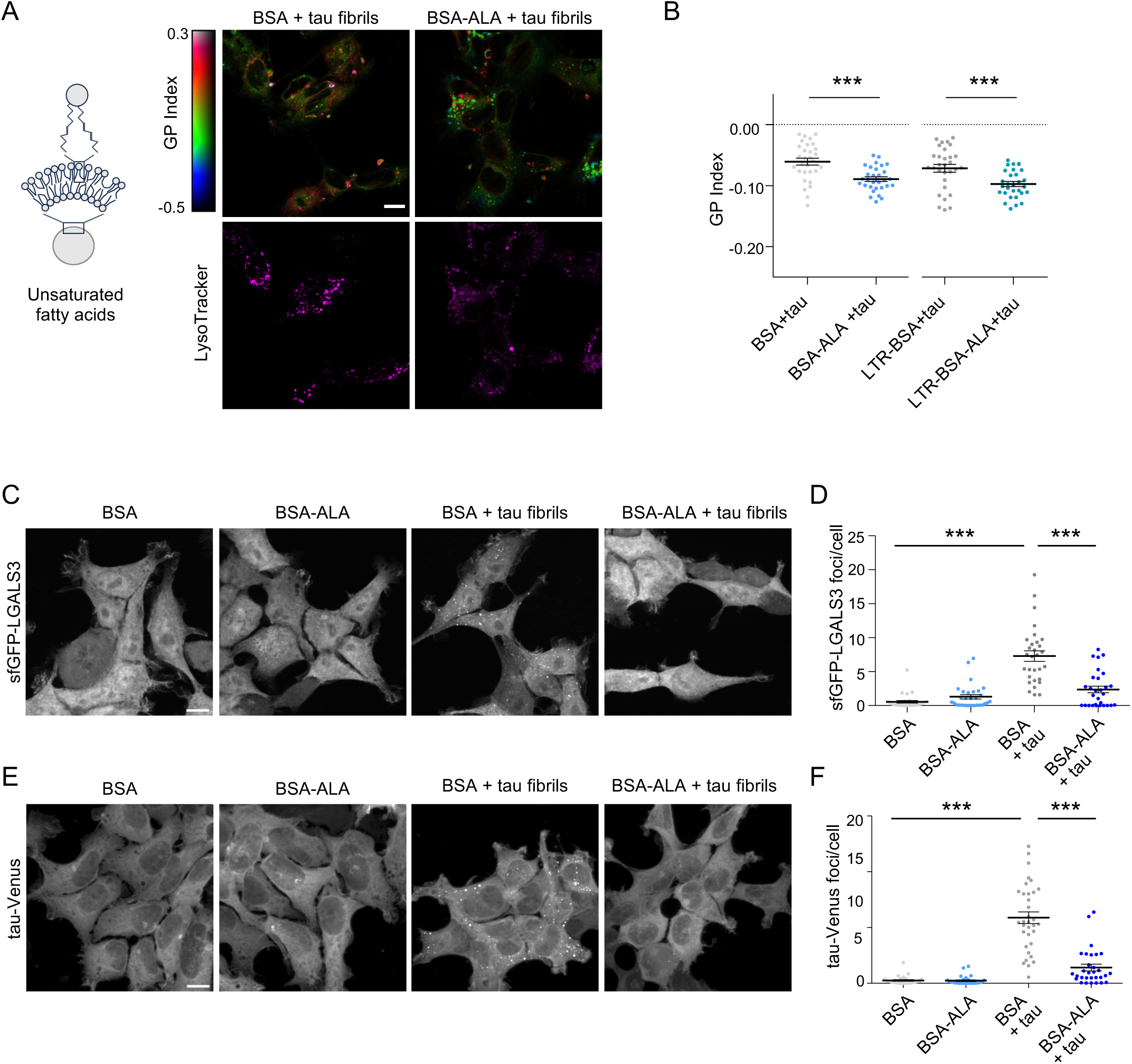
PUFA supplementation restores lysosomal membrane integrity and reduces seeded tau aggregation. **(A)** Left: unsaturated fatty acid membrane scheme. Right: Pseudo-colored images of SH-SY5Y cells pre-loaded with BSA control or 150 µM ALA conjugated to BSA (BSA-ALA) and treated with 1N4R tau fibrils showing the C-Laurdan GP Index at each pixel position. Lower panels show LysoTracker staining. Scale bar = 10 µm. **(B)** Quantification of GP values in SH-SY5Y cells pre-loaded with BSA control or 150 µM ALA conjugated to BSA (BSA-ALA) and exposed to 1N4R tau fibrils. GP values were measured across the whole cell (left) or restricted to LysoTracker-positive (LTR) regions (right). Statistical analysis was conducted using a two-way mixed-model ANOVA, followed by pairwise comparisons of estimated marginal means with Sidak correction for multiple comparisons. n= 3 independent experiments, with 10 images analyzed per experiment. **(C)** Maximum intensity projection of confocal z-stacks of HEK293T cells expressing sfGFP-LGALS3 pre-loaded with BSA or BSA-ALA with or without exposure to 1N4R tau fibrils. Scale bar = 10 µm. **(D)** Quantification of sfGFP-LGALS3 foci following indicated treatments. Statistical analysis comparing BSA + tau to other conditions was done using Kruskal-Wallis with Dunn’s post-hoc test. n = 3 independent experiments, with 10 images analyzed per experiment. *** = p < 0.001. **(E)** Maximum intensity projection of confocal z-stacks of tau-Venus biosensor cell line pre-loaded with BSA or BSA-ALA with or without exposure to 1N4R tau fibrils. Scale bar = 10 µm. **(F)** Quantification of tau-Venus foci following indicated treatments. Statistical analysis comparing BSA + tau to other conditions was done using Kruskal-Wallis with Dunn’s post-hoc test. n = 3 independent experiments, with 10 images analyzed per experiment. *** = p < 0.001.

Since ALA prevented endolysosomal rupture, we expected it to also prevent seeded tau aggregation. Indeed, tau-Venus cells pre-loaded with ALA exhibited significantly fewer foci when exposed to tau fibrils (Figure 5E, F and Figure S5F). Together, these data show that increasing lysosomal membrane fluidity counteracts tau fibril-induced endolysosomal membrane damage and limits downstream seeded tau aggregation.

Finally, we asked whether ALA would also reduce the toxicity associated with aggregated tau in our *C. elegans* model. Expression of F3ΔK281::mCherry in touch receptor neurons resulted in an age-dependent decline in the response to gentle touch from day 4 (first day of adulthood) to day 8 (day 5 of adulthood) (Figure 6A, and Figure S6)^20^. Supplementation of ALA significantly mitigated this behavioral deficit (Figure 6A and Figure S6) and reduced neurotoxicity (Figure 6B, C). Consistent with improved neuronal integrity, ALA treatment also reduced sfGFP::LGALS3 foci in touch receptor neurons (Figure 6D), indicating decreased endolysosomal damage in vivo. Together, these findings show that ALA-mediated increase in membrane fluidity reduces tau fibril-induced lysosomal membrane rigidification, endolysosomal damage, and seeded tau aggregation in human cells, while attenuating tau-associated neuronal dysfunction and endolysosomal damage in *C. elegans*.

**Figure 6.**
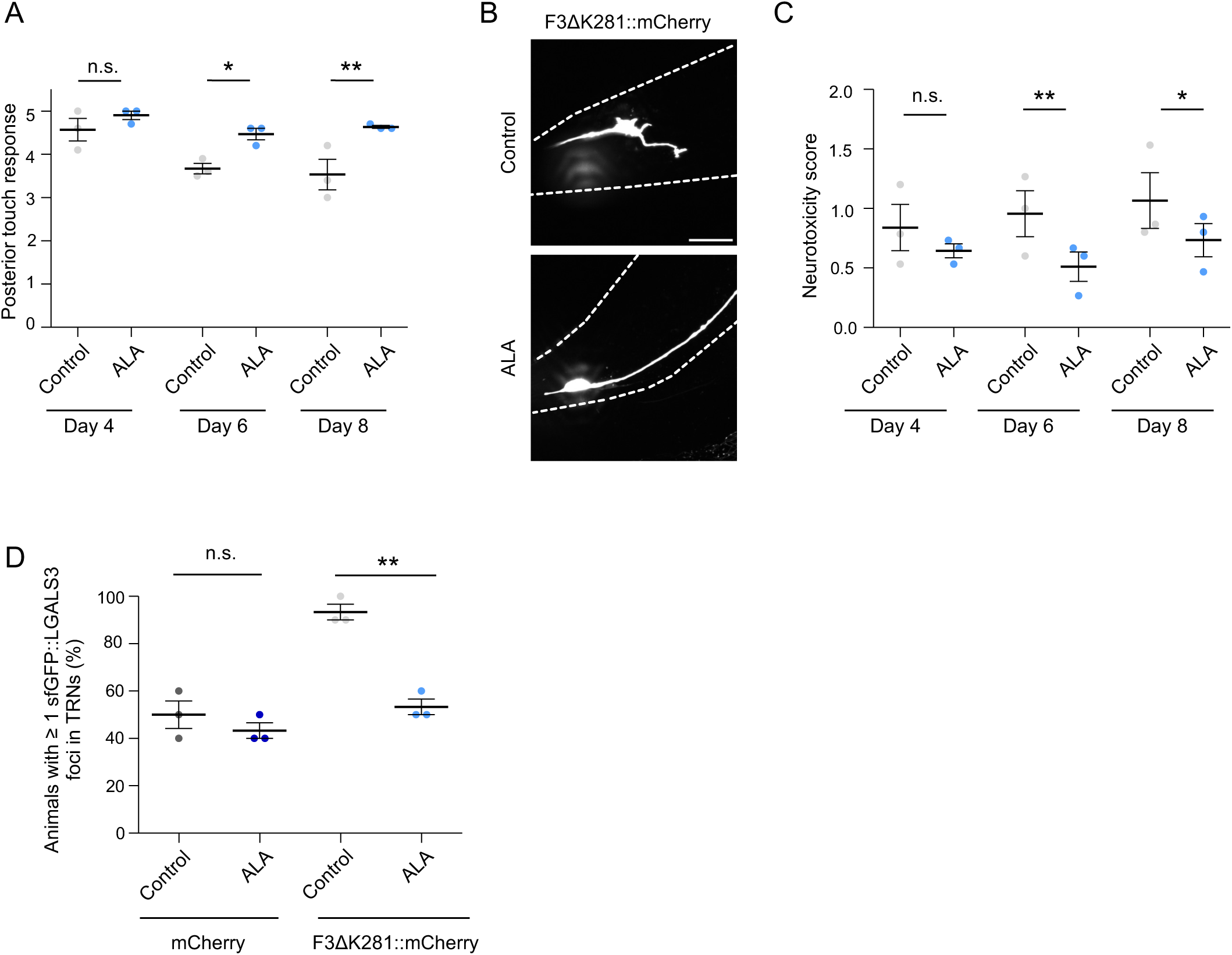
ALA improves neuronal function and reduces toxicity during aging. **(A)** Posterior touch response of animals expressing F3ΔK281::mCherry at indicated ages when grown on plates supplemented with ALA or ethanol solvent only control. Statistical analysis was done using two-way ANOVA with Bonferroni’s multiple comparison test. n = 3 independent experiments, with 10 animals analyzed per experiment. **(B)** Maximum intensity projection of confocal z-stacks of day 6 old animals expressing F3ΔK281::mCherry grown on EtOH solvent control or ALA plates. Scale bar = 20 µm. **(C)** Neurotoxicity score of PLM neurons of animals expressing F3ΔK281::mCherry grown on EtOH solvent control or ALA plates at indicated ages. Statistical analysis was done using two-way ANOVA with a Holm-Sidak’s multiple comparison test. n = 3 independent experiments, with 15 animals analyzed per experiment. **(D)** Quantification of the percentage of animals with ≥1 sfGFP::LGALS3 puncta in touch receptor neurons, upon co-expression of either mCherry control or F3ΔK281::mCherry and growth on EtOH solvent control or ALA-supplemented plates on day 5 (second day of adulthood). Statistical analysis was done using two-way ANOVA with Bonferroni’s multiple comparison test. Each dot represents an independent experiment, lines indicate the mean ± SEM. n= 3 independent experiments, with 10 animals analyzed per experiment. n.s.: not significant, * = p < 0.05, ** = p < 0.01.

## Discussion

Building on our previously published genome-wide screen in *C. elegans*, which identified SL metabolism as a regulator of endolysosomal integrity^20^, we investigated how genetic perturbation of enzymes involved in SL metabolism promotes endolysosomal membrane rupture and tau-associated phenotypes. We found that disruption of SL metabolism decreases endolysosomal membrane fluidity, making vesicles more prone to rupture. Fibrillar tau had a similar effect on membrane rigidity and acted additively with perturbation of SL metabolism to exacerbate endolysosomal damage. In human cell models, disruption of SL metabolism enhanced tau-fibril-induced rupture and seeded tau aggregation, whereas increasing membrane fluidity with ALA reduced tau-induced membrane rigidification, endolysosomal damage, and seeded aggregation. In *C. elegans*, ALA attenuated aggregated tau-associated neuronal dysfunction and endolysosomal damage. Together, these findings support a model in which disruption of SL metabolism and tau accumulation converge on endolysosomal membrane rigidification, thereby promoting endomembrane rupture, and allowing tau seeds to escape into the cytosol, where they seed the aggregation of soluble endogenously expressed tau.

Our results show that reducing the expression of genes involved in both, biosynthesis and degradation of SLs significantly reduced the fluidity of endolysosomal membranes, making them more fragile and prone to rupture. KD of the saposin *spp-10*, the sphingosine kinase *sphk-1*, and the sphingosine-1-phosphate lyase *spl-1* blocks the degradation of complex SLs and should lead to an increase in their relative abundance, and therefore to a decrease in membrane fluidity. KD of the very-long-chain (3R)-3-hydroxyacyl-CoA dehydratase *hpo-8*, which is involved in fatty acid elongation, has been demonstrated to promote the incorporation of long-chain UFAs into phospholipids, thereby restoring plasma membrane fluidity^39^. The absence of *hpo-8* leads to membrane rigidification^39^. Conversely, KD of the serine incorporator *R11H6.*2 and the 3-ketodihydrosphingosine reductase *Y37E11AM.3*, which function in the SL de novo synthesis pathway, should theoretically decrease total SL content and consequently increase membrane fluidity. The cytochrome b5 reductases *hpo-19* and *T05H4.4*, required for lipid desaturation, are involved in the conversion of dihydroceramide to ceramide and the biosynthesis of PUFAs^40^; KD of these genes is expected to reduce the levels of SLs and PUFAs, which could theoretically either increase or decrease membrane fluidity, depending on which metabolic pathway is predominantly affected.

Our observation that KD of all these genes results in decreased membrane fluidity is therefore seemingly counterintuitive. However, SLs form a complex metabolic network and can interconvert, allowing dynamic adaptation of SL levels. In addition, cells may compensate for the loss of certain SLs by altering the biosynthesis or turnover of other lipid molecules. This dynamic interplay makes it difficult to predict how individual gene knockdowns affect endolysosomal membrane composition and biophysical properties. Thus, genes that act at apparently opposing steps of SL metabolism could still converge on similar changes in membrane packing and fluidity. Lipidomic analyses will be important to determine how SL perturbations remodel cellular and organellar lipid composition. However, even detailed lipidomics would not by itself identify which lipid changes are responsible for the observed membrane rigidification. Indeed, membrane fluidity is not determined by a single lipid species, but by the combined properties of the entire membrane, including lipid abundance, saturation, acyl-chain length, head groups, sterol content, and membrane-associated proteins. Thus, an increase or decrease in a given lipid species cannot be directly translated into a predictable change in membrane fluidity without additional functional validation. Future lysosome-enriched or organelle-specific lipidomic approaches, ideally complemented by analyses of membrane-associated proteins, will therefore need to be combined with direct manipulation of candidate lipid or protein species, followed by measurements of membrane fluidity and rupture, to determine which molecular changes causally contribute to endolysosomal membrane rigidification.

Importantly, endolysosomal membrane rupture and global lysosomal function are related but not identical readouts. This distinction is supported by work showing that lipid dysregulation can induce lysosomal membrane permeabilization and lysosomal accumulation of endogenous protein aggregates without broadly impairing core lysosomal or proteasomal functions^41^. Thus, membrane damage can occur even when general lysosomal activity is not overtly disrupted. Conversely, a recent study independently identified SPHK-1 as an important regulator of lysosomal integrity in *C. elegans*, showing that strong *sphk-1* loss-of-function causes lysosomal sphingosine accumulation, membrane rupture, impaired degradative function, cargo accumulation, developmental defects, and reduced lifespan^42^. Together, these studies suggest that the functional consequences of disrupted SL metabolism can vary depending on allele strength, tissue, developmental stage, and assay conditions. In the present study, we focused on endolysosomal membrane fluidity and rupture because these membrane-level changes are directly linked to tau seed escape and seeded aggregation.

The central nervous system (CNS) is particularly rich in lipids. Approximately 50% of the brain’s dry weight consists of lipids with high concentrations of cholesterol, phospholipids and sphingolipids^43, 44^. Alterations in lipid and SL metabolism have been observed not only during the aging process, but also in various late-onset neurodegenerative diseases including AD^26, 45^. While aging is the major risk factor for AD, one of the most significant genetic risk factors for AD is the apolipoprotein E (ApoE) polymorphism, specifically the ApoE4 allele^46^. ApoE is the primary lipoprotein in the brain, crucial for lipid transport and metabolism. Besides cholesterol, the ApoE4 variant has been shown to influence SL metabolism, thereby affecting the pathogenesis of AD from its early stages^26, 45, 47^. Thus, although disturbances in SL homeostasis are increasingly recognized in AD, the mechanisms by which they contribute to disease progression remain poorly understood.

AD brains exhibit elevated levels of sphingosine, ceramide 1-phosphate, and ceramide compared to control subjects^45^. Multivariate analysis and machine learning also identified these SL species as key contributors to AD^45^. However, these analyses were performed in different brain regions without cell-type or subcellular resolution. We found that perturbation of SL metabolism primarily affected endolysosomal membranes, with little effect on plasma membrane fluidity in *C. elegans*. In human cells, LysoTracker-based analysis further revealed pronounced increases in GP values within lysosome-associated compartments. The lysosomal membrane is particularly rich in sphingolipids^48^, which may make it especially sensitive to perturbation of SL metabolism. Studies in yeast have shown that increased membrane lipid saturation affects the vacuole—the functional counterpart of mammalian lysosomes—as well as the nuclear envelope, whereas the plasma membrane and mitochondria remained relatively unaffected^49^. These findings indicate that cellular membranes vary in their resilience to lipid saturation stress, potentially due to differences in lipid composition and protective mechanisms^49, 50^. To better understand the implications of SL alterations in AD, it would be important to examine whether the observed changes primarily originate from endolysosomal membranes or also from other organelles, such as the endoplasmic reticulum and the nucleus, or from the plasma membrane.

We observed that fibrillar tau also increased membrane rigidity and exacerbated endolysosomal rupture. This suggests that tau accumulation and disruption of SL metabolism may converge on a common biophysical mechanism. This interpretation is consistent with recent ultrastructural studies showing that intralysosomal amyloid assemblies can physically deform and rupture lysosomal membranes^51, 52^. Whether this mechanism is specific to tau or also applies to other amyloid assemblies remains to be determined. In *C. elegans*, tau transmission sensitized endolysosomal membranes to SL perturbation, especially under sub-saturating RNAi conditions. The enhanced rupture phenotype is unlikely to result from a direct effect of RNAi on neuronal F3ΔK281::mCherry expression, as *C. elegans* neurons are largely refractory to systemic RNAi under the conditions used here^53^. Consistent with this, *sphk-1* RNAi did not alter hypodermal F3ΔK281::mCherry levels, supporting the interpretation that SL perturbation increases endolysosomal membrane damage rather than tau transmission itself.

These findings also raise the possibility that alterations in SL metabolism are not only a cause but also a consequence of tau-associated pathology^26, 45^. Accumulation of misfolded tau in lysosomes could impair lysosomal clearance capacity and alter lipid turnover, leading to accumulation of membrane lipids or other biomolecules. In turn, impaired SL metabolism could further reduce endolysosomal membrane fluidity and promote rupture. Such a cycle, in which imbalances in proteostasis and lipostasis negatively influence each other, may contribute to progressive lysosomal dysfunction and neurodegeneration^54^.

Manipulating membrane fluidity independently of SL enzymes further supported a role for membrane biophysics in tau-induced endolysosomal damage. Increasing membrane rigidity with PA exacerbated tau-induced rupture and seeded aggregation, whereas increasing membrane fluidity with ALA reduced tau-induced lysosomal membrane rigidification, endolysosomal damage, and seeded aggregation in cells and attenuated tau-associated neuronal dysfunction in *C. elegans*.

Given the high levels of saturated and trans fats in the Western diet^55^, it is plausible that dietary factors contribute to the dysregulation of lipid homeostasis in the brain. In line with a broader role of lipid saturation in AD, a recent plasma lipidomics study reported reduced highly unsaturated lipid species and increased saturated lipid species in women with AD^56^. The Mediterranean UFA-rich diet has been associated with healthy aging, while reduced UFA brain levels have been correlated with increased tau and Aβ pathology and decreased cognitive function in AD^57, 58^. Our findings therefore provide a possible mechanistic explanation for the beneficial effects of PUFA-rich diets or PUFA supplementation reported in AD-related contexts^59, 60^. However, PUFAs can also influence lipid signaling, oxidative stress responses, and broader membrane remodeling. We therefore cannot exclude additional direct or indirect effects of ALA. Nevertheless, the opposing effects of PA and ALA, together with the SL knockdown data, support the interpretation that membrane fluidity is a major determinant of endolysosomal membrane integrity and rupture in our models.

In sum, our study identifies endolysosomal membrane fluidity as a key determinant of vesicle integrity in the context of disrupted SL metabolism and tau accumulation. We propose that perturbation of SL metabolism and fibrillar tau converge on membrane rigidification, thereby reducing endolysosomal membrane integrity and promoting vesicle rupture, tau seed escape, and seeded tau aggregation. Restoring membrane fluidity may therefore represent a strategy to limit tau-induced endolysosomal damage and tau-associated toxicity, although the lipid species, membrane compartments, and additional cellular pathways contributing to this protection remain to be defined.

## Methods

### Maintenance of *C. elegans,* RNAi experiments and age synchronization

All animals were cultured using standard methods^61^. If not otherwise indicated, worms were grown on nematode growth medium (NGM) plates seeded with *E. coli* strain OP50 at 20 °C. For RNAi, NGM medium was supplemented with Ampicillin, Tetracycline, and IPTG, seeded with the respective HT115 *E. coli* RNAi clones (from the Ahringer RNAi library, for details see Table S2) and grown at 20 °C. For diluted RNAi experiments, RNAi was diluted with EV control bacteria. Animals were age-synchronized by bleaching. Briefly, gravid adults were dissolved in 20% sodium hypochlorite solution. The surviving *C. elegans* embryos were hatched overnight in M9 buffer with gentle rocking at 20 °C. The next day, appropriate amounts of L1 larvae were added to the plates.

### Cloning and generation of the neuronal sfGFP::LGALS3 strain

The sfGFP::LGALS3 reporter sequence was amplified from a previously cloned plasmid^20^ via PCR using suitable primers for subsequent Gibson assembly. The pCFJ150 vector backbone already containing an rgef-1p::GFP1-10::tbb2-3’UTR construct was cut via restriction digestion using NheI (NEB #R0131) and BglII (NEB **#**R0144) to remove the GFP1-10 sequence. The sfGFP::LGALS3 construct was subsequently cloned into this vector backbone in a Gibson assembly reaction using 0.08 pmol of plasmid backbone and a 5-fold molar excess of insert. 10 µl of fragment-vector mix was added to 10 µl Gibson assembly reaction mix containing 1.3x ISO buffer (6.5% w:v PEG-8000 [Sigma-Aldrich 89510], 130 mM Tris-HCl (Carl Roth, 4855.2) pH = 7.5, 13 mM MgCl2 (Carl Roth, A537.1), 13 mM DTT (Sigma-Aldrich 43816), 0.26 mM each dATP, dTTP, dCTP, and dGTP (NEB, N0446), 1.3 mM NAD (NEB, B9007S)), 0.05 U T5 exonuclease [NEB, M0663], 0.3 U Phusion polymerase (Thermo Fisher Scientific, F530), and 50 U Taq ligase [NEB, M0208L] and incubated at 50 °C for 2h. Subsequently the whole reaction was used to transform chemically competent *E. coli*. The extrachromosomal array was created by microinjecting the expression plasmid rgef-1p::sfGFP::LGALS3::tbb2-3’UTR (30 ng/µl) into the gonads of young adult wildtype *C. elegans* (N2s), together with a co-injection marker (15 ng/µl) for expression of RFP in coelomocytes. All strains generated or used in this study are found in Table S1.

### *C. elegans* fatty acid supplementation

Polyunsaturated fatty acids were dissolved in EtOH and stored in the dark at -20 °C under a N_2_ atmosphere to prevent oxidation. NGM was cooled after autoclaving to 55 °C. To enhance fatty acid distribution, NP-40 substitute (0.001% v/v) was added to the media and subsequently either 0.3 mM of the desired fatty acid solution or the ethanol solvent as control and stirred for 5 min before plate pouring^62^. Palmitic acid (PA) was supplemented by enriching the OP50 bacterial food source^63^. Overnight cultures were grown in the presence of 2 mM of PA dissolved in EtOH or EtOH alone. The next day, OP50 bacteria were pelleted by centrifugation (10 min at 4500x g) and washed 2x with M9 buffer^64^. 10x concentrated bacteria (in M9) were then seeded on NGM plates.

### Quantification of lysosomal rupture (galectin puncta assay) in *C. elegans*

Quantification of animals showing hypodermal lysosomal rupture was done as previously described^29, 65^. Briefly, age-synchronized animals expressing the sfGFP::LGALS3 construct in the hypodermis were seeded on OP50 or RNAi NGM plates. Scoring was done on day 5 (second day of adulthood) using a Leica M205 FA widefield binocular microscope. 20-50 worms were analyzed per replicate. *Sphk-1* mutant animals were obtained from CGC (CZ24969; *sphk-1(ju831)*) and crossed with animals expressing sfGFP::LGALS3 in the hypodermis.

Formation of sfGFP::LGALS3 foci in touch receptor neurons was quantified in day 5 (second day of adulthood) old animals co-expressing either mCherry or F3ΔK281::mCherry in touch receptor neurons using an Olympus IXplore SpinSR confocal microscope equipped with a UPlanSApo 60×/1.30 silicone oil objective. Foci formation was checked using a 488 nm laser at 80% power with an exposure time of 200 ms. An animal was counted as positive for endolysosomal rupture if it had at least one sfGFP:: LGALS3 foci in the PLM or ALM neurons.

### Mounting of live animals for imaging

Synchronized animals were mounted on 8-10% agarose in M9 buffer pads with a drop of mounting mix (2% (w/v) levamisole and 50% (v/v) nanosphere size standards (Thermo Fisher)) and covered with a coverslip.

### Fluorescence recovery after photobleaching (FRAP)

FRAP measurements to estimate membrane fluidity were done following a published protocol^66^. Animals were synchronized by bleaching and grown until day 5 (second day of adulthood) on the indicated RNAi or empty vector (EV) control plates. Directly before the FRAP experiments, animals were mounted at the Zeiss LSM 780 confocal microscope. FRAP measurements of either GFP enriched in the intestinal plasma membrane, or LAAT-1::mCherry located in intestinal or hypodermal lysosomal membranes were taken with 40X Water immersion objective. For the lysosomal membrane, mCherry-positive membranes were photobleached over a circular area (seven pixel radius) using 10 iterations of the 561 nm DPSS laser with 100% laser power transmission. Images were collected at a 12-bit intensity resolution over 128 × 128 pixels (digital zoom 6X) using a pixel dwell time of 3.15 μsec with 2% laser power transmission. For the plasma membrane, GFP-positive membranes were photobleached over a circular area (10 pixel radius) using 20 iterations of the 488 nm Argon laser with 100% laser power transmission. Images were collected at a 12-bit intensity resolution over 128 × 128 pixels (digital zoom 6X) using a pixel dwell time of 3.15 μsec with 10% laser power transmission. At least 6 pre-bleaching images were collected before the region of interest was bleached. The recovery of fluorescence was traced for 25s. Fluorescence recovery and T_half_ were calculated as previously described in Svensk et al.^67^.

### Quantification of spreading

Transmission of F3ΔK281::mCherry was quantified as previously published in Sandhof et al.^20^. In short, the hypodermis of age-synchronized worms expressing F3ΔK281::mCherry in touch receptor neurons were imaged on day 5 (second day of adulthood) using an Olympus IXplore SpinSR confocal microscope equipped with a UPlanSApo 60×/1.30 silicone oil objective. Images were acquired using a 561 nm laser at 30% power with an exposure time of 450 ms. Z-stacks were collected at 0.35 μm intervals, and hypodermal fluorescence intensity (integrated density) was quantified using ImageJ.

### Touch sensory assay and neurotoxicity in touch receptor neurons

Touch sensory assay was done as previously described^68^. Briefly, touch sensitivity was tested by gently stroking the tip of an eyebrow hair attached to a toothpick transversely across the anterior or posterior half of an animal. A touch-sensitive animal was one that stopped/moved away from the stimulus. 10 worms were examined for each strain in 3 biological replicates with 5 strokes in the anterior and posterior. For neurotoxicity scoring of the PLM neurons, live animals were imaged using an Olympus IXplore SpinSR Confocal microscope with an UplanS Apo 60x/1.30 Silicon oil objective. The neurotoxicity score was determined by adding a score of 1 for each of the following phenotypes: soma outgrowth, process branching, process break, wavy process. 15 animals were examined at each day of age in 3 independent biological replicates.

### Cell culture

SH-SY5Y and HEK293T cells were cultured in DMEM containing high glucose, GlutaMAX Supplement, pyruvate (Gibco), and 10% FBS (Gibco) at 37 °C and 5% CO_2_. Regular mycoplasma tests were performed (GATC Biotech). HEK293T cells expressing Venus-tagged full-length P301S mutant 0N4R tau (0N4R tauP301S-Venus) were kindly provided by Dr. William A. McEwan, Cambridge University^37^.

### Quantification of lysosomal rupture (galectin puncta assay) and tau-Venus foci in cells

Cells were imaged on an Olympus IXplore SpinSR Confocal microscope with an UplanS Apo 60x/1.30 Silicon oil objective. To detect the number of foci in an image, a difference of gaussian blur was applied to maximum projections to isolate foci from cytosolic sfGFP or Venus signal, respectively. Thresholding was applied using ‘moments’ threshold on FIJI. Number of foci was counted using the ‘analyze particles’ function. Cell number was counted manually to obtain number of foci/cell for each image.

### C-Laurdan dye measurement of membrane fluidity in worms and cells

Live worms were stained with 10 mM C-Laurdan dye (6-dodecanoyl-2-dimethylaminonaphthalene) (Thermo Fisher Scientific) as previously described^69^. ROI was drawn around the soma of the PLM using the mCherry signal and only the GP values in this ROI were collected. Cells were also stained with C-Laurdan at 15 µM for 2 hours in combination with LysoTracker (Thermo Fisher Scientific) at 50 nM, cells were then fixed in 4% PFA in PBS. Images were acquired with an DMI6000 confocal microscope and Leica software with a 63x oil-immersion objective, with a zoom factor of 4 or with a Leica SP8X WLL microscope, equipped with 405 nm laser, WLL2 laser (470 - 670 nm) and acusto-optical beam splitter. Images were acquired with a 63×1.4 objective. Samples were excited with a 405 nm laser and the emission recorded between 400 and 460 nm (ordered phase) and between 470 and 530 nm (disordered phase). Quantitative assessment of the membrane order was achieved by calculating the ratiometric relationship of the fluorescence intensity recorded in two spectral channels (GP value) by using an automated ImageJ macro, according to published guidelines^35^. LysoTracker-positive regions were identified by thresholding the LysoTracker channel, and GP values were extracted from these regions using the macro cited above.

### Knockdown of target genes by RNAi in cells

SPHK2 or a scrambled control siRNA were purchased from Thermo Scientific Dharmacon (ONTarget plus siRNA and Control Pool). Cells were seeded in DMEM with 10% FBS. siRNA was diluted in siRNA buffer (Final concentration 20 nM) before mixing with Lipofectamine 2000 for 20 min prior to being added to cells. To minimize Lipofectamine impact on the endolysosomal system, media was exchanged after 6 hours.

### SDS-PAGE and immunoblotting

Cells were pelleted by centrifugation followed by lysis in lysis buffer (10mM Tris, 100mM NaCl, 0.2% Triton X-100, 10mM EDTA) on ice for 20 min. The lysates were transferred into fresh Eppendorf tubes and centrifuged (1000 g for 1 min at 4 °C) in a tabletop centrifuge to remove cellular debris. The protein concentration was determined using protein assay dye reagent concentrate (Bio-Rad). Proteins were separated under denaturing conditions by SDS–PAGE and transferred onto a PVDF membrane (Carl Roth) by standard wet blotting protocols. Samples were probed with rabbit polyclonal anti-SPHK2 (1:5000, 9C5E1, Santa Cruz) primary antibody. Mouse monoclonal anti-GAPDH antibody (1:5000, clone GAPDH-71.1, Sigma-Aldrich) was used as loading control. HRP-conjugated anti-mouse and anti-rabbit IgG secondary antibodies (Bio-Rad) were used for subsequent ECL-based detection (Bio-Rad).

### Fatty acid treatment in cells

For α-linolenic acid (ALA), a stock solution was made in EtOH and stored in the dark at -20°C under a N_2_ atmosphere to prevent oxidation^70^. Palmitic acid (PA) was dissolved in 0.1 M NaOH according to Cousin et al.^71^. ALA and PA were then conjugated to BSA in 10% fatty acid free solution by shaking for 1 hour at 37°C to a concentration of 6 mM for ALA and 5 mM for PA^72^. Prior to preloading cells with BSA-PA, the solution was heated to 65 °C for 15 min. For assays with C-Laurdan, cells were preloaded with FAs for 3 hours before the addition of C-Laurdan for another 2 hours. If preloading was followed by seeding with tau, preloading with FAs was done for 3 hours before fibrillar tau was added for an additional 5 hours. C-Laurdan was spiked in for the last 2 hours. No media changes were done between steps. For experiments with HEK293T tau-Venus cells and HEK293T sfGFP-LGALS3 cells, preloading with FAs was done for 3 hours before seeding with 1N4R tau fibrils. Cells were then fixed after 48 hours.

### Monomeric and fibrillar tau

Full-length human 1N4R tau was expressed and purified as previously described^73^. The purified monomeric tau was dialyzed against PBS buffer containing 1 mM DTT, flash frozen in liquid nitrogen and stored at -80 °C. The concentration of monomeric tau was determined spectrophotometrically using an extinction coefficient at 280 nm of 7575 M^−1^ cm^−1^. Monomeric 1N4R tau was assembled into fibrils at 40 μM in the presence of 10 μM heparin, in PBS buffer containing 1 mM DTT under continuous shaking (600 rpm) for 5 days at 37 °C in an Eppendorf Thermomixer. The quality of 1N4R tau fibrils produced after 5 days was assessed by ThioflavinT binding, sedimentation, and transmission electron microscopy (TEM). The resulting 1N4R tau fibrils were spun at 25 °C and 75 000 g for 30 min. The pelleted fibrils were resuspended in PBS buffer with 1 mM DTT at 50 μM equivalent monomeric tau concentration. 1N4R tau fibrils were labeled by incubation with the 2 molar equivalents of lysine-reactive ATTO 550 (ATTO-TEC, GMBH) for 1 h at room temperature. The unreacted fluorophore was removed by two cycles of centrifugation at 75 000 g for 10 min and resuspension of the pellet in PBS. 1N4R tau fibrils were fragmented by sonication for 5 min in 2-ml Eppendorf tubes in a Vial Tweeter powered by an ultrasonic processor UIS250v (250 W, 2.4 kHz; Hielscher Ultrasonic, Teltow, Germany) to generate fibrillar particles with an average size 50 nm as assessed by TEM analysis. For seeding experiments, either monomeric or fibrillar tau was added directly into the cell culture media to a final concentration of 400 nM. For C-Laurdan experiments, tau was added to cells for 5h with the last 2h C-Laurdan and LysoTracker were spiked into the media. For lysosomal rupture and tau seeding, cells were fixed after 48 hours with tau.

### Statistical analysis

Statistical analysis was performed with either GraphPad Prism (GraphPad Software, Version 6h and 10.1.1) or in R (version 4.3.1, with packages ‘nlme’ version 3.1.166, ‘lme4’ version 1.1-35.5, ‘emmeans’ version 1.10.6, ‘dplyr’ version 1.1.4). Data were assessed for normal distribution by Shapiro Normality Test where appropriate. Parametric or non-parametric tests were applied as indicated in the figure legends. Data presentation, sample size (number of replicates, and sample size per experiment and condition), and the applied statistical tests are indicated in the figure legends for each experiment. For all cell culture data, data have been collected in 3 independent experiments with 10 images analyzed in each replicate. For C-Laurdan and LysoTracker, the data was analyzed in R using a two-way mixed-model ANOVA to account for non-independence of measurements acquired from the same experiment and/or image. Models were fitted using the lme4 package. Pairwise comparisons between conditions were performed on estimated marginal means using the emmeans package with Sidak correction for multiple comparisons. Significance levels: non-significant (n.s.) p > 0.05, ∗p ≤ 0.05, ∗∗p ≤ 0.01, and ∗∗∗p ≤ 0.001.

## Acknowledgements

We gratefully acknowledge the technical support of Sarah Wübbel. We extend our gratitude to Dr. Sunil Yeruva for his assistance with the use of the DMI6000 confocal microscope. We appreciate the help of Dr. Hana Nuskova and Dr. Aurelio Teleman (DKFZ, Heidelberg) with the fatty acid conjugation protocol. We thank Dr. Tamara Mikeladze-Dvali (LMU Munich) for her help with generating transgenic animals and for generously providing access to her microinjection system. The tauP301S-Venus expressing HEK293T cell line was kindly provided by Dr. William A. McEwan (Cambridge University)^37^. The pLenti PGK Puro DEST (w529-2) vector was a gift from Dr. Eric Campeau and Dr. Paul Kaufman^74^. We also thank Dr. Bin Liu and Dr. Marja Jäättelä (University of Copenhagen) for sharing the strain BIJ34 and a pPD49.26 expression plasmid coding for sfGFP::LGALS3 and Dr. Xiaochen Wang (Chinese Academy of Sciences, Beijing) for nematodes expressing the qxIs352 transgene. Some strains were provided by the Caenorhabditis Genetics Center (CGC), which is funded by the NIH Office of Research Infrastructure Programs (P40 OD010440).

## Funding

This work was supported by the Alzheimer Forschungs Initiative (AFI, grant #21053) (to CNK) and The Fondation pour la Recherche Médicale, contract ALZ201912009776 (to RM).

## Author contributions

Conceptualization J.T., C.A.S., C.N.K.

Formal Analysis J.T., C.A.S., N.M.

Funding acquisition R.M., C.N.K.

Investigation J.T., C.A.S., N.M., D.E.

Methodology J.T., C.A.S., N.M., S.B.N.N

Project administration C.N.K.

Resources S.B.N.N.

Supervision R.M., C.N.K.

Visualization J.T., C.N.K.

Writing – original draft J.T., C.A.S., C.N.K.

Writing – review & editing J.T., C.A.S., N.M., D.E., R.M., C.N.K.

## Declaration of interests

The authors declare no competing interest.

## Supplemental information

Figures S1–S6, Tables S1-S2

## Data availability

All data supporting the findings of this study are available within the paper and its supplementary information. *C. elegans* strains and plasmids generated in this study are available upon request.

## Notes

### Competing Interest Statement

The authors have declared no competing interest.

### Summary of Updates

We have strengthened the lysosomal specificity of the membrane-fluidity measurements by combining C-Laurdan imaging with LysoTracker-based analysis, expanded the human cell data by including SH-SY5Y neuroblastoma cells, added genetic validation using a sphk-1 mutant strain, clarified the C. elegans strains used in this study, corrected the interpretation of Lipofectamine-associated effects, and revised the Discussion to better address lipidomics, lysosomal function, PUFA pleiotropy, and the relationship between membrane rupture and tau seeding. We also adjusted the wording throughout the manuscript to more precisely distinguish our experimental interventions from model-based interpretations.

## References

(1) Clavaguera, F.; Bolmont, T.; Crowther, R. A.; Abramowski, D.; Frank, S.; Probst, A.; Fraser, G.; Stalder, A. K.; Beibel, M.; Staufenbiel, M.;, et al. Transmission and spreading of tauopathy in transgenic mouse brain. Nature cell biology 2009, 11 (7), 909–909. DOI: 10.1038/NCB1901.

(2) Walker, L. C.; Jucker, M. The prion principle and Alzheimer’s disease. Science 2024, 385 (6715), 1278–1279. DOI: 10.1126/SCIENCE.ADQ5252.

(3) Biel, D.; Brendel, M.; Rubinski, A.; Buerger, K.; Janowitz, D.; Dichgans, M.; Franzmeier, N.; Alzheimer’s Disease Neuroimaging, I. Tau-PET and in vivo Braak-staging as prognostic markers of future cognitive decline in cognitively normal to demented individuals. Alzheimer’s research & therapy 2021, 13 (1), 137. DOI: 10.1186/s13195-021-00880-x From NLM Medline.

(4) Frost, B.; Jacks, R. L.; Diamond, M. I. Propagation of tau misfolding from the outside to the inside of a cell. J Biol Chem 2009, 284 (19), 12845–12852. DOI: 10.1074/jbc.M808759200.

(5) Sanders, D. W.; Kaufman, S. K.; DeVos, S. L.; Sharma, A. M.; Mirbaha, H.; Li, A.; Barker, S. J.; Foley, A. C.; Thorpe, J. R.; Serpell, L. C.;, et al. Distinct tau prion strains propagate in cells and mice and define different tauopathies. Neuron 2014, 82 (6), 1271–1288. DOI: 10.1016/j.neuron.2014.04.047.

(6) Shrivastava, A. N.; Redeker, V.; Pieri, L.; Bousset, L.; Renner, M.; Madiona, K.; Mailhes-Hamon, C.; Coens, A.; Buee, L.; Hantraye, P.;, et al. Clustering of Tau fibrils impairs the synaptic composition of alpha3-Na(+)/K(+)-ATPase and AMPA receptors. EMBO J 2019, 38 (3). DOI: 10.15252/embj.201899871.

(7) Caballero, B.; Wang, Y.; Diaz, A.; Tasset, I.; Juste, Y. R.; Stiller, B.; Mandelkow, E. M.; Mandelkow, E.; Cuervo, A. M. Interplay of pathogenic forms of human tau with different autophagic pathways. Aging cell 2018, 17 (1). DOI: 10.1111/acel.12692 From NLM Medline.

(8) Menzies, F. M.; Fleming, A.; Rubinsztein, D. C. Compromised autophagy and neurodegenerative diseases. Nat Rev Neurosci 2015, 16 (6), 345–357. DOI: 10.1038/nrn3961 From NLM Medline.

(9) Nixon, R. A.; Wegiel, J.; Kumar, A.; Yu, W. H.; Peterhoff, C.; Cataldo, A.; Cuervo, A. M. Extensive involvement of autophagy in Alzheimer disease: an immuno-electron microscopy study. J Neuropathol Exp Neurol 2005, 64 (2), 113–122.

(10) Hou, W. C.; Massey, L. A.; Rhoades, D.; Wu, Y.; Ren, W.; Frank, C.; Overkleeft, H. S.; Kelly, J. W. A PIKfyve modulator combined with an integrated stress response inhibitor to treat lysosomal storage diseases. Proc Natl Acad Sci U S A 2024, 121 (34), e2320257121. DOI: 10.1073/pnas.2320257121 From NLM Medline.

(11) Lim, S. H. Y.; Hansen, M.; Kumsta, C. Molecular Mechanisms of Autophagy Decline during Aging. Cells 2024, 13 (16). DOI: 10.3390/cells13161364 From NLM Medline.

(12) Caballero, B.; Bourdenx, M.; Luengo, E.; Diaz, A.; Sohn, P. D.; Chen, X.; Wang, C.; Juste, Y. R.; Wegmann, S.; Patel, B.;, et al. Acetylated tau inhibits chaperone-mediated autophagy and promotes tau pathology propagation in mice. Nature communications 2021, 12 (1), 2238. DOI: 10.1038/s41467-021-22501-9 From NLM Medline.

(13) Chen, J. J.; Nathaniel, D. L.; Raghavan, P.; Nelson, M.; Tian, R.; Tse, E.; Hong, J. Y.; See, S. K.; Mok, S. A.; Hein, M. Y.;, et al. Compromised function of the ESCRT pathway promotes endolysosomal escape of tau seeds and propagation of tau aggregation. J Biol Chem 2019. DOI: 10.1074/jbc.RA119.009432.

(14) Flavin, W. P.; Bousset, L.; Green, Z. C.; Chu, Y.; Skarpathiotis, S.; Chaney, M. J.; Kordower, J. H.; Melki, R.; Campbell, E. M. Endocytic vesicle rupture is a conserved mechanism of cellular invasion by amyloid proteins. Acta Neuropathol 2017. DOI: 10.1007/s00401-017-1722-x.

(15) Tuck, B. J.; Miller, L. V. C.; Katsinelos, T.; Smith, A. E.; Wilson, E. L.; Keeling, S.; Cheng, S.; Vaysburd, M. J.; Knox, C.; Tredgett, L.;, et al. Cholesterol determines the cytosolic entry and seeded aggregation of tau. Cell reports 2022, 39 (5), 110776. DOI: 10.1016/j.celrep.2022.110776 From NLM Medline.

(16) Dimou, E.; Katsinelos, T.; Meisl, G.; Tuck, B. J.; Keeling, S.; Smith, A. E.; Hidari, E.; Lam, J. Y. L.; Burke, M.; Lovestam, S.;, et al. Super-resolution imaging unveils the self-replication of tau aggregates upon seeding. Cell reports 2023, 42 (7), 112725. DOI: 10.1016/j.celrep.2023.112725 From NLM Medline.

(17) Polanco, J. C.; Hand, G. R.; Briner, A.; Li, C.; Götz, J. Exosomes induce endolysosomal permeabilization as a gateway by which exosomal tau seeds escape into the cytosol. Acta Neuropathol 2021, 141 (2), 235–256. DOI: 10.1007/s00401-020-02254-3 From NLM.

(18) Rose, K.; Jepson, T.; Shukla, S.; Maya-Romero, A.; Kampmann, M.; Xu, K.; Hurley, J. H. Tau fibrils induce nanoscale membrane damage and nucleate cytosolic tau at lysosomes. Proc Natl Acad Sci U S A 2024, 121 (22), e2315690121. DOI: 10.1073/pnas.2315690121 From NLM Medline.

(19) Kroemer, G.; Jaattela, M. Lysosomes and autophagy in cell death control. Nature reviews. Cancer 2005, 5 (11), 886–897. DOI: 10.1038/nrc1738.

(20) Sandhof, C. A.; Martin, N.; Tittelmeier, J.; Schlueter, A.; Pezzali, M.; Schoendorf, D. C.; Lange, T.; Reinhardt, P.; Ried, J. S.; Liang, S.;, et al. A novel C. elegans model for MAPT/Tau spreading reveals genes critical for endolysosomal integrity and seeded MAPT/Tau aggregation. Autophagy 2025. DOI: 10.1080/15548627.2025.2551676 From NLM Publisher.

(21) Breslow, D. K.; Weissman, J. S. Membranes in balance: mechanisms of sphingolipid homeostasis. Mol Cell 2010, 40 (2), 267–279. DOI: 10.1016/j.molcel.2010.10.005 From NLM Medline.

(22) Jimenez-Rojo, N.; Leonetti, M. D.; Zoni, V.; Colom, A.; Feng, S.; Iyengar, N. R.; Matile, S.; Roux, A.; Vanni, S.; Weissman, J. S.;, et al. Conserved Functions of Ether Lipids and Sphingolipids in the Early Secretory Pathway. Curr Biol 2020, 30 (19), 3775–3787 e3777. DOI: 10.1016/j.cub.2020.07.059 From NLM Medline.

(23) Gault, C. R.; Obeid, L. M.; Hannun, Y. A. An overview of sphingolipid metabolism: from synthesis to breakdown. Adv Exp Med Biol 2010, 688, 1–23. DOI: 10.1007/978-1-4419-6741-1_1 From NLM Medline.

(24) Futerman, A. H.; Hannun, Y. A. The complex life of simple sphingolipids. EMBO Rep 2004, 5 (8), 777–782. DOI: 10.1038/sj.embor.7400208 From NLM Medline.

(25) Czubowicz, K.; Jesko, H.; Wencel, P.; Lukiw, W. J.; Strosznajder, R. P. The Role of Ceramide and Sphingosine-1-Phosphate in Alzheimer’s Disease and Other Neurodegenerative Disorders. Mol Neurobiol 2019, 56 (8), 5436–5455. DOI: 10.1007/s12035-018-1448-3 From NLM Medline.

(26) van Echten-Deckert, G.; Walter, J. Sphingolipids: critical players in Alzheimer’s disease. Prog Lipid Res 2012, 51 (4), 378–393. DOI: 10.1016/j.plipres.2012.07.001 From NLM Medline.

(27) Huotari, J.; Helenius, A. Endosome maturation. EMBO J 2011, 30 (17), 3481–3500. DOI: 10.1038/emboj.2011.286 From NLM Medline.

(28) Yong, J.; Villalta, J. E.; Vu, N.; Kukurugya, M. A.; Olsson, N.; Lopez, M. P.; Lazzari-Dean, J. R.; Hake, K.; McAllister, F. E.; Bennett, B. D.;, et al. Impairment of lipid homeostasis causes lysosomal accumulation of endogenous protein aggregates through ESCRT disruption. eLife 2024, 12. DOI: 10.7554/eLife.86194 From NLM Medline.

(29) Aits, S.; Kricker, J.; Liu, B.; Ellegaard, A. M.; Hamalisto, S.; Tvingsholm, S.; Corcelle-Termeau, E.; Hogh, S.; Farkas, T.; Holm Jonassen, A.;, et al. Sensitive detection of lysosomal membrane permeabilization by lysosomal galectin puncta assay. Autophagy 2015, 11 (8), 1408–1424. DOI: 10.1080/15548627.2015.1063871.

(30) Hanel, V.; Pendleton, C.; Witting, M. The sphingolipidome of the model organism Caenorhabditis elegans. Chem Phys Lipids 2019, 222, 15–22. DOI: 10.1016/j.chemphyslip.2019.04.009 From NLM Medline.

(31) Cheng, X.; Jiang, X.; Tam, K. Y.; Li, G.; Zheng, J.; Zhang, H. Sphingolipidomic Analysis of C. elegans reveals Development- and Environment-dependent Metabolic Features. Int J Biol Sci 2019, 15 (13), 2897–2910. DOI: 10.7150/ijbs.30499 From NLM Medline.

(32) D’Auria, L.; Bongarzone, E. R. Fluid levity of the cell: Role of membrane lipid architecture in genetic sphingolipidoses. J Neurosci Res 2016, 94 (11), 1019–1024. DOI: 10.1002/jnr.23750 From NLM Medline.

(33) Liu, B.; Du, H.; Rutkowski, R.; Gartner, A.; Wang, X. LAAT-1 is the lysosomal lysine/arginine transporter that maintains amino acid homeostasis. Science 2012, 337 (6092), 351–354. DOI: 10.1126/science.1220281.

(34) Morck, C.; Olsen, L.; Kurth, C.; Persson, A.; Storm, N. J.; Svensson, E.; Jansson, J. O.; Hellqvist, M.; Enejder, A.; Faergeman, N. J.;, et al. Statins inhibit protein lipidation and induce the unfolded protein response in the non-sterol producing nematode Caenorhabditis elegans. Proc Natl Acad Sci U S A 2009, 106 (43), 18285–18290. DOI: 10.1073/pnas.0907117106 From NLM Medline.

(35) Owen, D. M.; Rentero, C.; Magenau, A.; Abu-Siniyeh, A.; Gaus, K. Quantitative imaging of membrane lipid order in cells and organisms. Nat Protoc 2011, 7 (1), 24–35. DOI: 10.1038/nprot.2011.419 From NLM Medline.

(36) Barucha-Kraszewska, J.; Kraszewski, S.; Ramseyer, C. Will C-Laurdan dethrone Laurdan in fluorescent solvent relaxation techniques for lipid membrane studies? Langmuir 2013, 29 (4), 1174–1182. DOI: 10.1021/la304235r From NLM Medline.

(37) McEwan, W. A.; Falcon, B.; Vaysburd, M.; Clift, D.; Oblak, A. L.; Ghetti, B.; Goedert, M.; James, L. C. Cytosolic Fc receptor TRIM21 inhibits seeded tau aggregation. Proc Natl Acad Sci U S A 2017, 114 (3), 574–579. DOI: 10.1073/pnas.1607215114 From NLM Medline.

(38) Nachman, E.; Wentink, A. S.; Madiona, K.; Bousset, L.; Katsinelos, T.; Allinson, K.; Kampinga, H.; McEwan, W. A.; Jahn, T. R.; Melki, R.;, et al. Disassembly of Tau fibrils by the human Hsp70 disaggregation machinery generates small seeding-competent species. J Biol Chem 2020, 295 (28), 9676–9690. DOI: 10.1074/jbc.RA120.013478.

(39) Ruiz, M.; Devkota, R.; Kaper, D.; Ruhanen, H.; Busayavalasa, K.; Radovic, U.; Henricsson, M.; Kakela, R.; Boren, J.; Pilon, M. AdipoR2 recruits protein interactors to promote fatty acid elongation and membrane fluidity. J Biol Chem 2023, 299 (6), 104799. DOI: 10.1016/j.jbc.2023.104799 From NLM Medline.

(40) Zhang, Y.; Wang, H.; Zhang, J.; Hu, Y.; Zhang, L.; Wu, X.; Su, X.; Li, T.; Zou, X.; Liang, B. The cytochrome b5 reductase HPO-19 is required for biosynthesis of polyunsaturated fatty acids in Caenorhabditis elegans. Biochim Biophys Acta 2016, 1861 (4), 310–319. DOI: 10.1016/j.bbalip.2016.01.009 From NLM Medline.

(41) Yong, J.; Villalta, J. E.; Vu, N.; Kukurugya, M. A.; Olsson, N.; López, M. P.; Lazzari-Dean, J. R.; Hake, K.; McAllister, F. E.; Bennett, B. D.;, et al. Impairment of lipid homeostasis causes lysosomal accumulation of endogenous protein aggregates through ESCRT disruption. eLife 2024, 12. DOI: 10.7554/ELIFE.86194.

(42) Li, Y.; Zhang, J.; Li, M.; Yang, L.; Wang, X. Sphingosine kinase SPHK-1 maintains sphingolipid metabolism to protect lysosome membrane integrity in C. elegans. Mol Biol Cell 2026, 37 (1), ar1. DOI: 10.1091/mbc.E25-04-0182 From NLM Medline.

(43) Yoon, J. H.; Seo, Y.; Jo, Y. S.; Lee, S.; Cho, E.; Cazenave-Gassiot, A.; Shin, Y. S.; Moon, M. H.; An, H. J.; Wenk, M. R.;, et al. Brain lipidomics: From functional landscape to clinical significance. Sci Adv 2022, 8 (37), eadc9317. DOI: 10.1126/sciadv.adc9317 From NLM Medline.

(44) Hornemann, T. Mini review: Lipids in Peripheral Nerve Disorders. Neurosci Lett 2021, 740, 135455. DOI: 10.1016/j.neulet.2020.135455 From NLM Medline.

(45) Uranbileg, B.; Isago, H.; Sakai, E.; Kubota, M.; Saito, Y.; Kurano, M. Alzheimer’s disease manifests abnormal sphingolipid metabolism. Front Aging Neurosci 2024, 16, 1368839. DOI: 10.3389/fnagi.2024.1368839 From NLM PubMed-not-MEDLINE.

(46) Corder, E. H.; Saunders, A. M.; Strittmatter, W. J.; Schmechel, D. E.; Gaskell, P. C.; Small, G. W.; Roses, A. D.; Haines, J. L.; Pericak-Vance, M. A. Gene dose of apolipoprotein E type 4 allele and the risk of Alzheimer’s disease in late onset families. Science 1993, 261 (5123), 921–923. DOI: 10.1126/science.8346443 From NLM Medline.

(47) Kosicek, M.; Zetterberg, H.; Andreasen, N.; Peter-Katalinic, J.; Hecimovic, S. Elevated cerebrospinal fluid sphingomyelin levels in prodromal Alzheimer’s disease. Neurosci Lett 2012, 516 (2), 302–305. DOI: 10.1016/j.neulet.2012.04.019 From NLM Medline.

(48) Fabri, J.; de Sa, N. P.; Malavazi, I.; Del Poeta, M. The dynamics and role of sphingolipids in eukaryotic organisms upon thermal adaptation. Prog Lipid Res 2020, 80, 101063. DOI: 10.1016/j.plipres.2020.101063 From NLM Medline.

(49) Romanauska, A.; Kohler, A. Lipid saturation controls nuclear envelope function. Nat Cell Biol 2023, 25 (9), 1290–1302. DOI: 10.1038/s41556-023-01207-8 From NLM Medline.

(50) Harayama, T.; Riezman, H. Understanding the diversity of membrane lipid composition. Nat Rev Mol Cell Biol 2018, 19 (5), 281–296. DOI: 10.1038/nrm.2017.138 From NLM Medline.

(51) Elias, R. D.; O’Neill, R. T.; Demiralp, II; Allen, S.; Serwas, D.; Siems, H.; Montabana, E. A.; Ash, C.; Yacoubian, D. A.; Lederberg, O. L.; et al. Cathepsin C-Catalyzed Ligation Generates Intralysosomal Amyloid Fibrils from Dipeptide Esters. bioRxiv 2025. DOI: 10.64898/2025.12.23.696283 From NLM PubMed-not-MEDLINE.

(52) Li, D.; Zhang, W.; Medina, M.; Stuke, J. F. M.; Schwarz, A.; Brill, J.; Brenner, J.; Kraus, F.; Ohlerich, S.; Lizarrondo, J.; et al. Cathepsin-dependent amyloid formation drives mechanical rupture of lysosomal membranes. bioRxiv 2026. DOI: 10.64898/2026.01.17.700056 From NLM PubMed-not-MEDLINE.

(53) Calixto, A.; Chelur, D.; Topalidou, I.; Chen, X.; Chalfie, M. Enhanced neuronal RNAi in C. elegans using SID-1. Nat Methods 2010, 7 (7), 554–559. DOI: 10.1038/nmeth.1463 From NLM Medline.

(54) Tittelmeier, J.; Nussbaum-Krammer, C. Broken Balance: Emerging Cross-Talk Between Proteostasis and Lipostasis in Neurodegenerative Diseases. Cells 2025, 14 (11). DOI: 10.3390/cells14110845 From NLM Medline.

(55) Rakhra, V.; Galappaththy, S. L.; Bulchandani, S.; Cabandugama, P. K. Obesity and the Western Diet: How We Got Here. Mo Med 2020, 117 (6), 536–538. From NLM Medline.

(56) Wretlind, A.; Xu, J.; Chen, W.; Velayudhan, L.; Ashton, N. J.; Zetterberg, H.; Proitsi, P.; Legido-Quigley, C. Lipid profiling reveals unsaturated lipid reduction in women with Alzheimer’s disease. Alzheimers Dement 2025, 21 (8), e70512. DOI: 10.1002/alz.70512 From NLM Medline.

(57) Snowden, S. G.; Ebshiana, A. A.; Hye, A.; An, Y.; Pletnikova, O.; O’Brien, R.; Troncoso, J.; Legido-Quigley, C.; Thambisetty, M. Association between fatty acid metabolism in the brain and Alzheimer disease neuropathology and cognitive performance: A nontargeted metabolomic study. PLoS Med 2017, 14 (3), e1002266. DOI: 10.1371/journal.pmed.1002266 From NLM Medline.

(58) Bischoff-Ferrari, H. A.; Gangler, S.; Wieczorek, M.; Belsky, D. W.; Ryan, J.; Kressig, R. W.; Stahelin, H. B.; Theiler, R.; Dawson-Hughes, B.; Rizzoli, R.;, et al. Individual and additive effects of vitamin D, omega-3 and exercise on DNA methylation clocks of biological aging in older adults from the DO-HEALTH trial. Nat Aging 2025. DOI: 10.1038/s43587-024-00793-y From NLM Publisher.

(59) Shinto, L.; Quinn, J.; Montine, T.; Dodge, H. H.; Woodward, W.; Baldauf-Wagner, S.; Waichunas, D.; Bumgarner, L.; Bourdette, D.; Silbert, L.;, et al. A randomized placebo-controlled pilot trial of omega-3 fatty acids and alpha lipoic acid in Alzheimer’s disease. J Alzheimers Dis 2014, 38 (1), 111–120. DOI: 10.3233/JAD-130722 From NLM Medline.

(60) Wood, A. H. R.; Chappell, H. F.; Zulyniak, M. A. Dietary and supplemental long-chain omega-3 fatty acids as moderators of cognitive impairment and Alzheimer’s disease. Eur J Nutr 2022, 61 (2), 589–604. DOI: 10.1007/s00394-021-02655-4 From NLM Medline.

(61) Brenner, S. The genetics of Caenorhabditis elegans. Genetics 1974, 77 (1), 71–94.

(62) Svensk, E.; Stahlman, M.; Andersson, C. H.; Johansson, M.; Boren, J.; Pilon, M. PAQR-2 regulates fatty acid desaturation during cold adaptation in C. elegans. PLoS Genet 2013, 9 (9), e1003801. DOI: 10.1371/journal.pgen.1003801 From NLM Medline.

(63) Ruiz, M.; Bodhicharla, R.; Svensk, E.; Devkota, R.; Busayavalasa, K.; Palmgren, H.; Stahlman, M.; Boren, J.; Pilon, M. Membrane fluidity is regulated by the C. elegans transmembrane protein FLD-1 and its human homologs TLCD1/2. Elife 2018, 7. DOI: 10.7554/eLife.40686 From NLM Medline.

(64) Devkota, R.; Svensk, E.; Ruiz, M.; Stahlman, M.; Boren, J.; Pilon, M. The adiponectin receptor AdipoR2 and its Caenorhabditis elegans homolog PAQR-2 prevent membrane rigidification by exogenous saturated fatty acids. PLoS Genet 2017, 13 (9), e1007004. DOI: 10.1371/journal.pgen.1007004 From NLM Medline.

(65) Sandhof, C. A.; Hoppe, S. O.; Druffel-Augustin, S.; Gallrein, C.; Kirstein, J.; Voisine, C.; Nussbaum-Krammer, C. Reducing INS-IGF1 signaling protects against non-cell autonomous vesicle rupture caused by SNCA spreading. Autophagy 2020, 16 (5), 878–899. DOI: 10.1080/15548627.2019.1643657.

(66) Devkota, R.; Pilon, M. FRAP: A Powerful Method to Evaluate Membrane Fluidity in Caenorhabditis elegans. Bio Protoc 2018, 8 (13), e2913. DOI: 10.21769/BioProtoc.2913 From NLM PubMed-not-MEDLINE.

(67) Svensk, E.; Devkota, R.; Stahlman, M.; Ranji, P.; Rauthan, M.; Magnusson, F.; Hammarsten, S.; Johansson, M.; Boren, J.; Pilon, M. Caenorhabditis elegans PAQR-2 and IGLR-2 Protect against Glucose Toxicity by Modulating Membrane Lipid Composition. PLoS Genet 2016, 12 (4), e1005982. DOI: 10.1371/journal.pgen.1005982 From NLM Medline.

(68) Chalfie, M.; Sulston, J. E.; White, J. G.; Southgate, E.; Thomson, J. N.; Brenner, S. The neural circuit for touch sensitivity in Caenorhabditis elegans. J Neurosci 1985, 5 (4), 956–964.

(69) Jeong, J. H.; Han, J. S.; Jung, Y.; Lee, S. M.; Park, S. H.; Park, M.; Shin, M. G.; Kim, N.; Kang, M. S.; Kim, S.;, et al. A new AMPK isoform mediates glucose-restriction induced longevity non-cell autonomously by promoting membrane fluidity. Nature communications 2023, 14 (1), 288. DOI: 10.1038/s41467-023-35952-z From NLM Medline.

(70) Oliveira, A. F.; Cunha, D. A.; Ladriere, L.; Igoillo-Esteve, M.; Bugliani, M.; Marchetti, P.; Cnop, M. In vitro use of free fatty acids bound to albumin: A comparison of protocols. Biotechniques 2015, 58 (5), 228–233. DOI: 10.2144/000114285 From NLM Medline.

(71) Cousin, S. P.; Hugl, S. R.; Wrede, C. E.; Kajio, H.; Myers, M. G., Jr.; Rhodes, C. J. Free fatty acid-induced inhibition of glucose and insulin-like growth factor I-induced deoxyribonucleic acid synthesis in the pancreatic beta-cell line INS-1. Endocrinology 2001, 142 (1), 229–240. DOI: 10.1210/endo.142.1.7863 From NLM Medline.

(72) Nuskova, H.; Cortizo, F. G.; Schwenker, L. S.; Sachsenheimer, T.; Diakonov, E. E.; Tiebe, M.; Schneider, M.; Lohbeck, J.; Reid, C.; Kopp-Schneider, A.;, et al. Competition for cysteine acylation by C16:0 and C18:0 derived lipids is a global phenomenon in the proteome. J Biol Chem 2023, 299 (9), 105088. DOI: 10.1016/j.jbc.2023.105088 From NLM Medline.

(73) Tardivel, M.; Begard, S.; Bousset, L.; Dujardin, S.; Coens, A.; Melki, R.; Buee, L.; Colin, M. Tunneling nanotube (TNT)-mediated neuron-to neuron transfer of pathological Tau protein assemblies. Acta neuropathologica communications 2016, 4 (1), 117. DOI: 10.1186/s40478-016-0386-4.

(74) Campeau, E.; Ruhl, V. E.; Rodier, F.; Smith, C. L.; Rahmberg, B. L.; Fuss, J. O.; Campisi, J.; Yaswen, P.; Cooper, P. K.; Kaufman, P. D. A versatile viral system for expression and depletion of proteins in mammalian cells. PLoS One 2009, 4 (8), e6529. DOI: 10.1371/journal.pone.0006529.

(75) Chitwood, D. J.; Lusby, W. R.; Thompson, M. J.; Kochansky, J. P.; Howarth, O. W. The glycosylceramides of the nematode Caenorhabditis elegans contain an unusual, branched-chain sphingoid base. Lipids 1995, 30 (6), 567–573. DOI: 10.1007/BF02537032 From NLM Medline.

(76) Zhang, H.; Abraham, N.; Khan, L. A.; Hall, D. H.; Fleming, J. T.; Gobel, V. Apicobasal domain identities of expanding tubular membranes depend on glycosphingolipid biosynthesis. Nat Cell Biol 2011, 13 (10), 1189–1201. DOI: 10.1038/ncb2328 From NLM Medline.

(77) Zhu, H.; Shen, H.; Sewell, A. K.; Kniazeva, M.; Han, M. A novel sphingolipid-TORC1 pathway critically promotes postembryonic development in Caenorhabditis elegans. eLife 2013, 2, e00429. DOI: 10.7554/eLife.00429 From NLM Medline.

(78) Scholz, J.; Helmer, P. O.; Nicolai, M. M.; Bornhorst, J.; Hayen, H. Profiling of sphingolipids in Caenorhabditis elegans by two-dimensional multiple heart-cut liquid chromatography - mass spectrometry. J Chromatogr A 2021, 1655, 462481. DOI: 10.1016/j.chroma.2021.462481 From NLM Medline.

