## Supplemental Figures for "Disruption of sphingolipid metabolism promotes tau seeding through endolysosomal membrane rigidification and rupture"

#### Figure S1

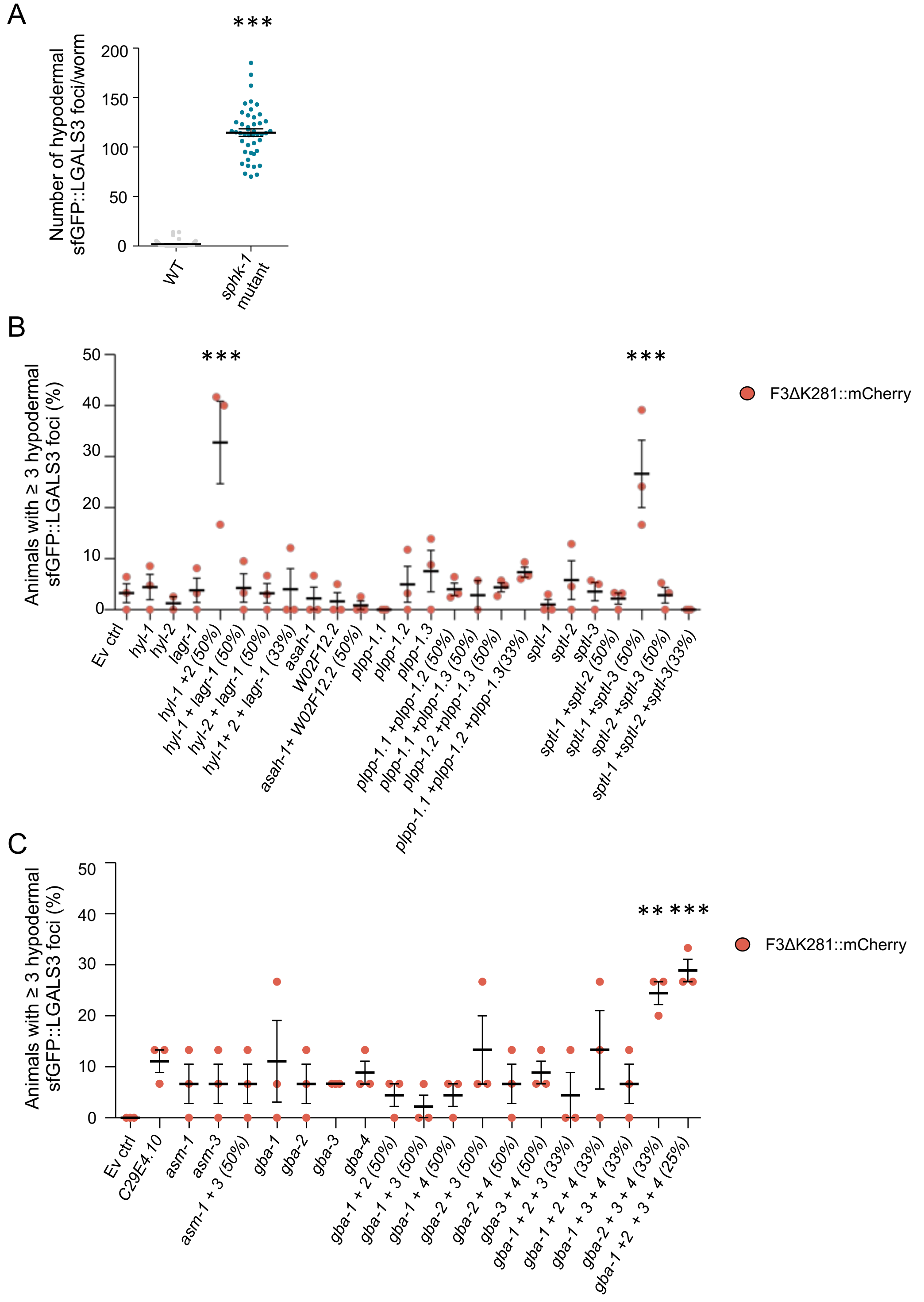

**Figure S1.** ***Sphk-1*** **and additional sphingolipid metabolic genes affect endolysosomal rupture.** **(A)** Genetic validation of the *sphk-1* RNAi phenotype using a *sphk-1* mutant. Quantification of sfGFP::LGALS3 foci per worm on day 5 (second day of adulthood) in wild-type (WT) and *sphk-1* mutant animals. Each dot represents one animal; data are shown as mean ± SEM. n=3 biological repeats with 15 animals per replicate. Statistical analysis was done using Student's t-test.

**(B, C)** Co-KD of redundant genes involved in SL metabolism identifies additional regulators of endolysosomal rupture. Quantification of the percentage of animals on day 5 (second day of adulthood) expressing F3ΔK281::mCherry in touch receptor neurons with ≥ 3 hypodermal sfGFP::LGALS3 foci, upon indicated single and co-KDs. Percentages shown next to gene names indicate the relative amount of each RNAi bacterial clone in co-KD conditions. (B) Among the genes involved in SL biosynthesis, co-KDs of *sptl-1* and *-3* and *hyl-1* and *-2* significantly induce endolysosomal rupture compared with the empty vector control. (C) For genes related to SL degradation, co-KD of *gba-2*, *-3*, and *-4* or *gba-1*, *-2*, *-3* and *-4* resulted in a significant increase in endolysosomal rupture. Data are shown as mean ± SEM. N = 45-60 animals from three biological replicates. Statistical analysis was done using one-way ANOVA with Dunnett's post-hoc test, ** = $p < 0.01$, *** = $p < 0.001$.

#### Figure S2

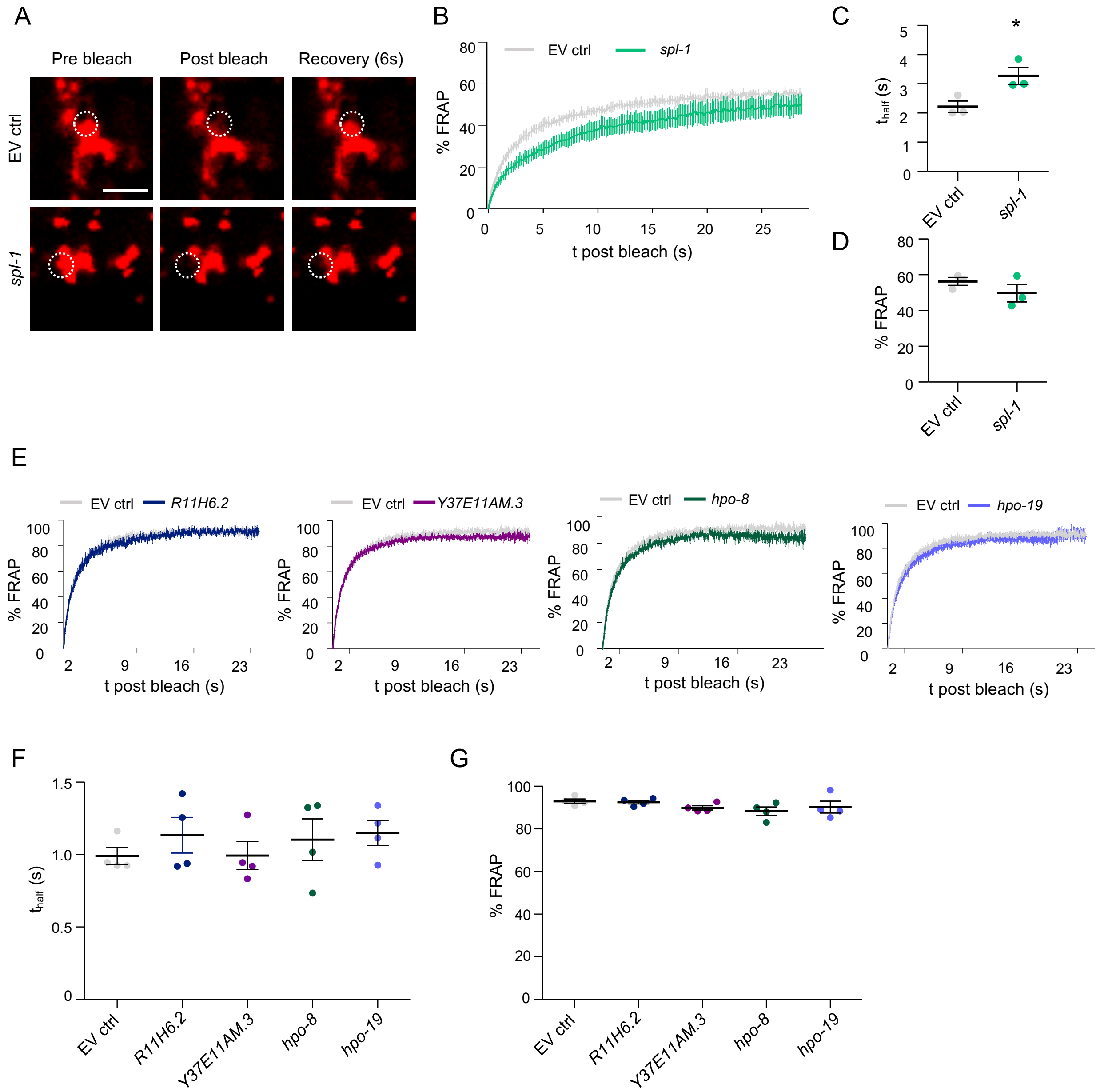


**Figure S2. Disruption of SL metabolism decreases lysosomal membrane fluidity.**

**(A)** Representative confocal single plane images from a FRAP experiment in animals on day 5 (second day of adulthood) expressing a mCherry tagged Lysosomal Lysine/Arginine Transporter 1 (LAAT-1::mCherry) in the intestine grown either on empty vector or *spl-1* RNAi plates. Dashed circles outline the bleach spots. Scale bar = 5 µm.

**(B)** Combined FRAP curves of LAAT-1::mCherry in intestinal lysosomes of animals on day 5 (second day of adulthood) grown either on empty vector or *spl-1* RNAi plates. Curves are normalized to the pre-bleach intensity as 100% and the first post-bleach intensity as 0%.

**(C, D)** Mean $t_{half}$ (C) and maximal % recoverable fluorescence values (D) calculated from the LAAT-1::mCherry FRAP experiments.

**(E)** FRAP curves of prenylated GFP enriched on the intestinal plasma membrane of animals on day 5 (second day of adulthood) grown either on empty vector or the indicated RNAi plates. Curves are normalized to the pre-bleach intensity set as 100% and the first post-bleach intensity as 0%.

**(F, G)** Mean $t_{half}$ (F) and maximal % recoverable fluorescence values (G) calculated from the GFP FRAP experiments. Data, including FRAP curves, represented as means ± SEM from 5 – 12 FRAP measurements on day 5 animals (second day of adulthood). Statistical analysis comparing RNAi conditions to the empty vector control was done using one-way ANOVA with Dunnett's post-hoc test. * = $p < 0.05$.

### Figure S3

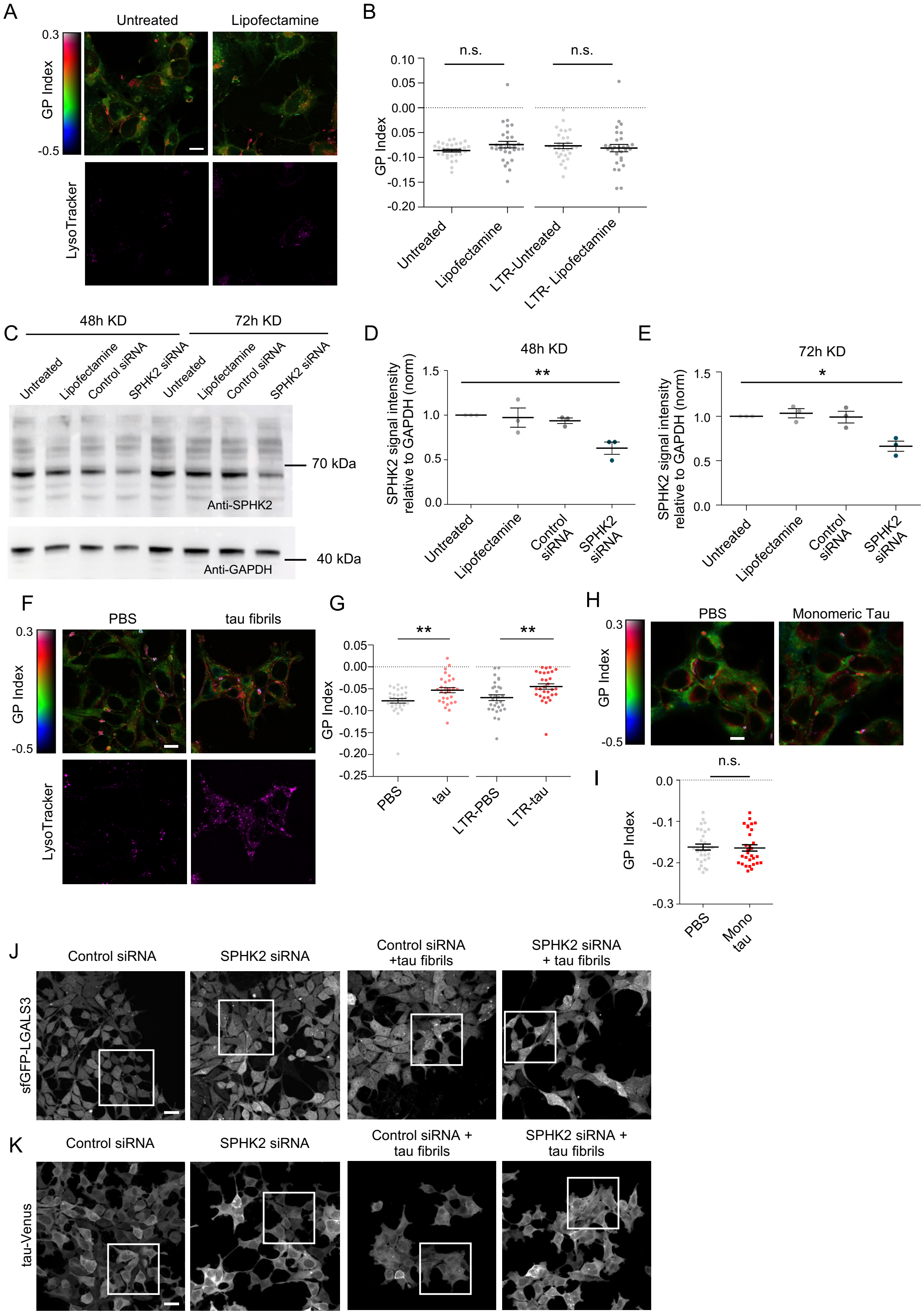

A
Untreated
Lipofectamine
0.3
GP Index
-0.5
LysoTracker
B
n.s.
n.s.
0.10
0.05
0.00
0.05
-0.10
-0.15
-0.20
GP Index
Untreated
Lipofectamine
LTR-Untreated
LTR- Lipofectamine
C
48h KD
72h KD
Untreated
Lipofectamine
Control siRNA
SPHK2 siRNA
Untreated
Lipofectamine
Control siRNA
SPHK2 siRNA
70 kDa
Anti-SPHK2
40 kDa
Anti-GAPDH
D
48h KD
**
SPHK2 signal intensity relative to GAPDH (norm)
Untreated
Lipofectamine
Control siRNA
SPHK2 siRNA
E
72h KD
*
SPHK2 signal intensity relative to GAPDH (norm)
Untreated
Lipofectamine
Control siRNA
SPHK2 siRNA
F
PBS
tau fibrils
0.3
GP Index
-0.5
LysoTracker
G
**
**
GP Index
PBS
tau
LTR-PBS
LTR-tau
H
PBS
Monomeric Tau
0.3
GP Index
-0.5
I
n.s.
GP Index
PBS
Mono tau
J
Control siRNA
SPHK2 siRNA
Control siRNA +tau fibrils
SPHK2 siRNA + tau fibrils
sfGFP-LGALS3
K
Control siRNA
SPHK2 siRNA
Control siRNA + tau fibrils
SPHK2 siRNA + tau fibrils
tau-Venus

**Figure S3. KD of SPHK2 and exposure to fibrillar tau decreases membrane fluidity in human cells.**

**(A)** Upper panels: pseudo-colored images of SH-SY5Y cells untreated or transfected with Lipofectamine showing the C-Laurdan GP Index. Lower panels: LysoTracker staining. Scale bar = 10 µm.

**(B)** Quantification of GP values in SH-SY5Y cells that were untreated or treated with Lipofectamine alone. GP values were measured either across the whole cell (left) or specifically within LysoTracker-positive (LTR) regions (right). No significant differences were observed (see also Figure 3C). Statistical analysis was performed using a two-way mixed-model ANOVA followed by pairwise comparisons of estimated marginal means with Sidak correction for multiple comparisons. n = 3 independent experiments, with 10 images analyzed per experiment.

**(C)** Western blot analysis of total cell lysates from SH-SY5Y cells after 48 h or 72 h of no treatment, Lipofectamine treatment alone, or transfection with control or SPHK2 siRNA using Lipofectamine. SPHK2 and GAPDH protein levels were assessed using anti-SPHK2 and anti-GAPDH antibodies, respectively (see methods section).

**(D, E)** Quantification of SPHK2 protein levels relative to GAPDH after 48 h (D) or 72 h (E) of indicated controls or SPHK2 siRNA treatment. Each dot represents an independent experiment, with lines indicating mean ± SEM. Statistical analysis was performed using one-way ANOVA followed by Dunnett's post hoc test.

**(F)** Upper panels: pseudo-colored images of HEK293T cells exposed to PBS control or 1N4R tau fibrils showing the C-Laurdan GP Index. Lower panels: LysoTracker staining. Scale bar = 10 µm.

**(G)** Quantification of GP values in control and HEK293T cells exposed to PBS control or 1N4R tau fibrils. Increased GP values indicate increased membrane rigidity following tau fibril treatment. Statistical analysis was performed using a two-way mixed-model ANOVA followed by pairwise comparisons of estimated marginal means with Sidak correction for multiple comparisons. n = 3 independent experiments, with 10 images analyzed per experiment.

**(H)** Pseudo-colored images of HEK293T cells exposed to PBS control or monomeric 1N4R tau displaying the C-Laurdan GP Index. Scale bar = 10 µm.

**(I)** Quantification of GP values in HEK293T cells exposed to PBS control or monomeric 1N4R tau. Statistical analysis was performed using Student's *t*-test. n = 3 independent experiments, with 10 images analyzed per experiment.

**(J)** Max. intensity projection of confocal z-stacks of HEK293T cells expressing sfGFP-LGALS3 following control or SPHK2 siRNA treatment, with or without 1N4R tau fibril seeding. White box indicates the zoomed in section depicted in Figure 3F. Scale bar = 20 µm.

**(K)** Max. intensity projection of confocal z-stacks of P301S mutant tau-Venus biosensor cell line following control or SPHK2 siRNA treatment, with or without 1N4R tau fibril seeding. White box indicates the zoomed in section depicted in Figure 3H. Scale bar = 20 µm. n.s.: not significant, * = $p < 0.05$, ** = $p < 0.01$.

### Figure S4

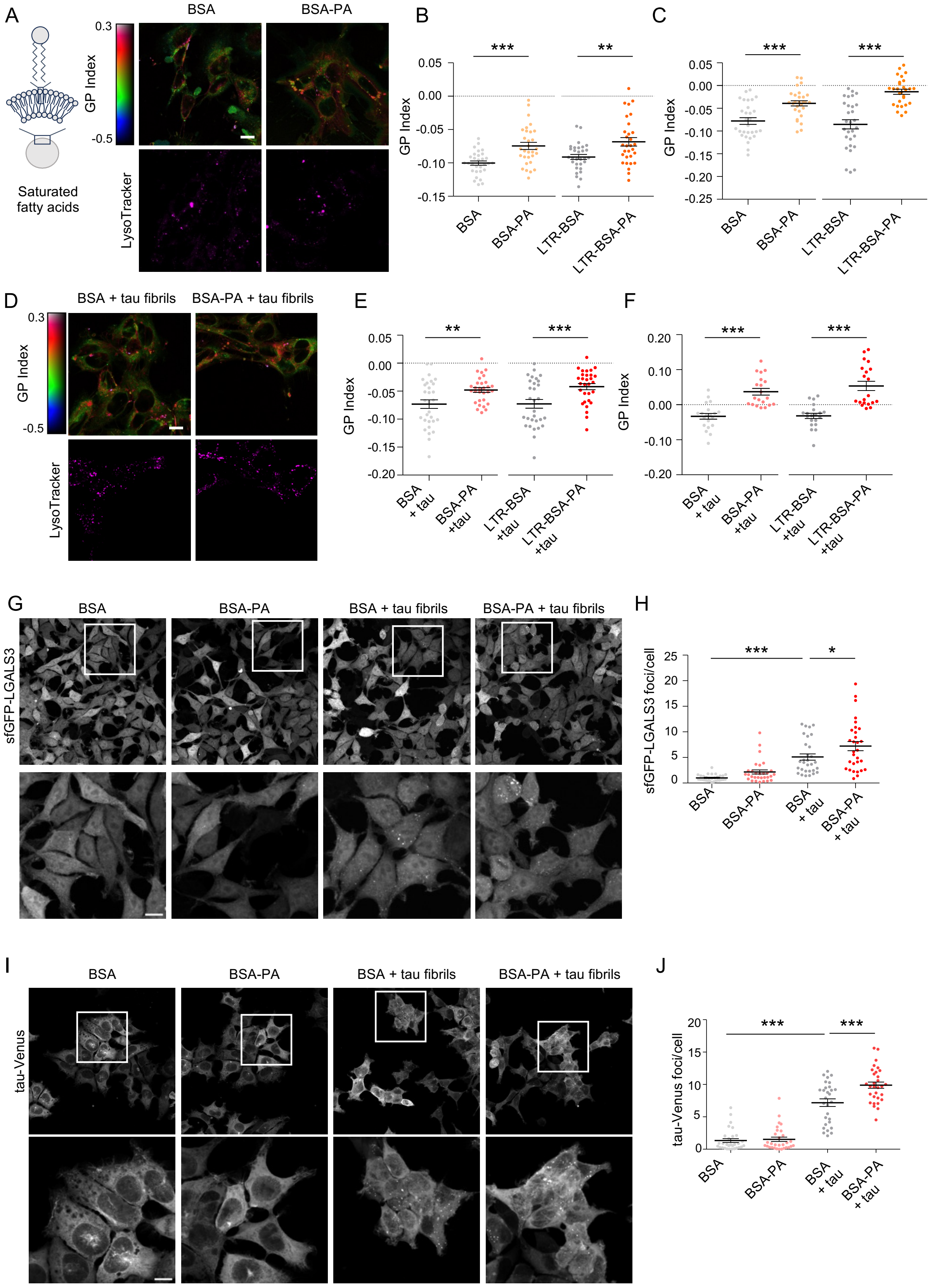

A
Saturated fatty acids
BSA
BSA-PA
GP Index
0.3
-0.5
LysoTracker
B
C
***
**
***
***
BSA
BSA-PA
LTR-BSA
LTR-BSA-PA
D
BSA + tau fibrils
BSA-PA + tau fibrils
E
F
BSA + tau
BSA-PA +tau
LTR-BSA +tau
LTR-BSA-PA +tau
G
H
sfGFP-LGALS3
sfGFP-LGALS3 foci/cell
*
I
J
tau-Venus
tau-Venus foci/cell

**Figure S4. Saturated fatty acids decrease membrane fluidity and exacerbate seeded tau aggregation.**

**(A)** Left: saturated fatty acid membrane scheme. Right: upper panels: pseudo-colored images of SH-SY5Y cells pre-loaded with BSA or 50 µM PA conjugated to BSA (BSA-PA) displaying the C-Laurdan GP Index. Lower panels: LysoTracker staining. Scale bar = 10 µm.

**(B)** Quantification of GP values in SH-SY5Y cells upon PA treatment. GP values were measured either across the whole cell (left) or specifically within LysoTracker-positive (LTR) regions (right). Statistical analysis was conducted using a two-way mixed-model ANOVA followed by pairwise comparisons of estimated marginal means with Sidak correction for multiple comparisons. n = 3 independent experiments, with 10 images analyzed per experiment.

**(C)** Quantification of GP values in HEK293T cells after PA treatment. GP values were measured either across the whole cell (left) or specifically within LysoTracker-positive (LTR) regions (right). Statistical analysis was performed using a two-way mixed-model ANOVA, followed by pairwise comparisons of estimated marginal means with Sidak correction for multiple comparisons. n = 3 independent experiments, with 10 images analyzed per experiment.

**(D)** Upper panels: Pseudo-colored images of SH-SY5Y cells pre-loaded with BSA or 50 µM PA conjugated to BSA (BSA-PA) and subsequently exposed to 1N4R tau fibrils displaying the C-Laurdan GP Index. Lower panels: LysoTracker staining. Scale bar = 10 µm.

**(E)** Quantification of GP values in SH-SY5Y cells pre-loaded with PA and subsequently exposed to 1N4R tau fibrils. GP values were measured either across the whole cell (left) or specifically within LysoTracker-positive (LTR) regions (right). Statistical analysis was conducted using a two-way mixed-model ANOVA, followed by pairwise comparisons of estimated marginal means with Sidak correction for multiple comparisons. n = 3 independent experiments, with 10 images analyzed per experiment.

**(F)** Quantification of GP values in HEK293T cells preloaded with PA and subsequently exposed to 1N4R tau fibrils. GP values were measured either across the whole cell (left) or specifically within LysoTracker-positive (LTR) regions (right). Statistical analysis was performed using a two-way mixed-model ANOVA, followed by pairwise comparisons of estimated marginal means with Sidak correction for multiple comparisons. n = 2 independent experiments, with 10 images analyzed per experiment.

**(G)** Max. intensity projection of confocal z-stacks of HEK293T cells expressing sfGFP-LGALS3 pre-loaded with BSA or BSA-PA and subsequently exposed to PBS control or 1N4R tau fibrils. White box indicates the zoomed in section depicted in lower panel. Scale bar = 10 μm.

**(H)** Quantification of sfGFP-LGALS3 foci in HEK293T cells pre-loaded with BSA or BSA-PA and subsequently exposed to PBS control or 1N4R tau fibrils. Statistical analysis comparing BSA + tau to other conditions was done using one-way ANOVA with Dunnett's post-hoc test. n = 3 independent experiments, with 10 images analyzed per experiment.

**(I)** Max. intensity projection of confocal z-stacks of tau biosensor cells expressing Venus-tagged full-length P301S tau pre-loaded with BSA or BSA-PA with or without exposure to 1N4R tau fibrils. White box indicates the zoomed in section depicted in lower panel. Scale bar = 10 μm.

**(J)** Quantification of P301S tau-Venus foci following indicated treatments. Statistical analysis comparing BSA + tau to other conditions was done using one-way ANOVA with Dunnett's post-hoc test. n = 3 independent experiments, 10 images analyzed per experiment. * = $p < 0.05$, ** = $p < 0.01$, *** = $p < 0.001$.

### Figure S5

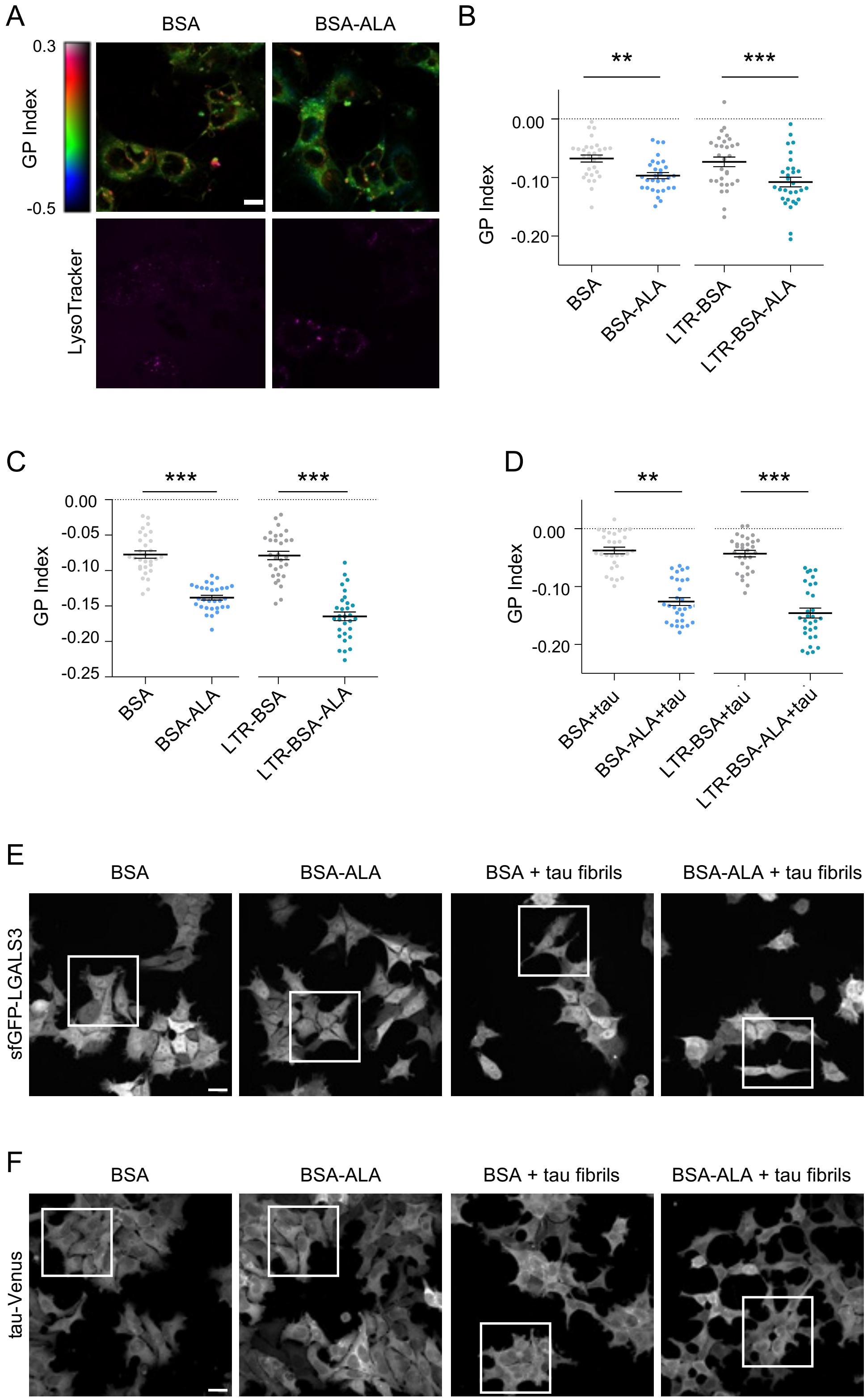
A
BSA
BSA-ALA
0.3
GP Index
-0.5
LysoTracker
B
**
***
0.00
-0.10
-0.20
GP Index
BSA
BSA-ALA
LTR-BSA
LTR-BSA-ALA
C
***
***
0.00
-0.05
-0.10
-0.15
-0.20
-0.25
GP Index
BSA
BSA-ALA
LTR-BSA
LTR-BSA-ALA
D
**
***
0.00
-0.10
-0.20
GP Index
BSA+tau
BSA-ALA+tau
LTR-BSA+tau
LTR-BSA-ALA+tau
E
BSA
BSA-ALA
BSA + tau fibrils
BSA-ALA + tau fibrils
sfGFP-LGALS3
F
BSA
BSA-ALA
BSA + tau fibrils
BSA-ALA + tau fibrils
tau-Venus

**Figure S5. PUFA supplementation restores lysosomal membrane integrity and reduces seeded tau aggregation.**

**(A)** Upper panels: Pseudo-colored images of SH-SY5Y cells pre-loaded with BSA control or 150 µM ALA conjugated to BSA (BSA-ALA) displaying the GP Index. Lower panels: LysoTracker staining. Scale bar = 10 µm.

**(B)** Quantification of GP values in SH-SY5Y cells preloaded with BSA or BSA-ALA. GP values were measured either across the whole cell or specifically within LysoTracker-positive (LTR) regions. Statistical analysis was conducted using a two-way mixed-model ANOVA with each experiment/image as random effects, with pairwise comparisons of estimated marginal means with Sidak correction for multiple comparisons. n = 3 independent experiments, 10 images each.

**(C, D)** Quantification of GP values in HEK293T cells preloaded with BSA or BSA-ALA and subsequently exposed to PBS control (C) or 1N4R tau fibrils (D). GP values were measured either across the whole cell or specifically within LysoTracker-positive (LTR) regions. Statistical analysis was conducted using a two-way mixed-model ANOVA followed by pairwise comparisons of estimated marginal means with Sidak correction for multiple comparisons. n = 3 independent experiments, with 10 images analyzed per experiment.

**(E)** Max intensity projection of confocal z-stacks of HEK293T cells expressing sfGFP-LGALS3 pre-loaded with BSA or BSA-ALA with or without exposure to 1N4R tau fibrils. White box indicates the zoomed in section depicted in Figure 5C. Scale bar = 20 µm.

**(F)** Max intensity projection of whole confocal z-stacks of P301S tau-Venus biosensor cell line pre-loaded with BSA or BSA-ALA with or without exposure to 1N4R tau fibrils. The white boxes delineate the zoomed images presented in Figure 5E. Scale bar = 20 µm. ** = $p < 0.01$, *** = $p < 0.001$.

**Figure S6**

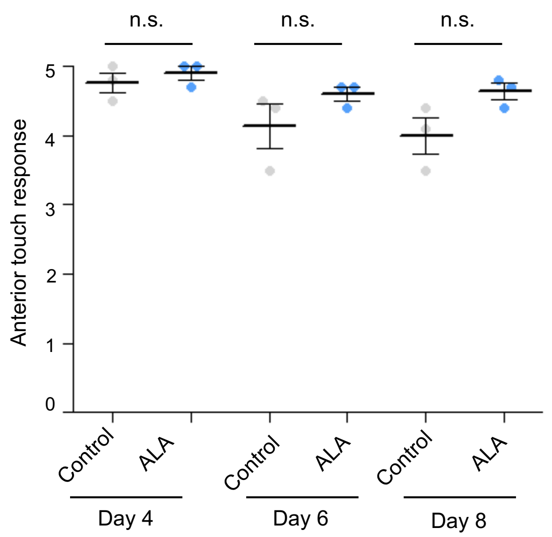


**Figure S6. The impact of ALA on anterior touch response.**

Anterior touch response of animals expressing F3ΔK281::mCherry at indicated ages when grown on plates supplemented with ALA or ethanol solvent only control. Statistical analysis was done using two-way ANOVA with Bonferroni's multiple comparison test. n = 3 independent experiments, with 10 worms analyzed per replicate. n.s.: not significant.
